# Genetic network analysis uncovers spatial variation in diversity and connectivity of a species presenting a continuous distribution

**DOI:** 10.1101/2024.01.22.576731

**Authors:** Cory Fournier, Micheline Manseau, Bridget Redquest, Leon Andrew, Allicia Kelly, Dave Hervieux, Troy Hegel, Gigi Pittoello, Vicki Trim, Dennis Brannen, Paul Wilson

**Affiliations:** Landscape Science and Technology Division, Environment and Climate Change Canada, Ottawa, Ontario, Canada; Biology Department, Trent University, Peterborough, Ontario, Canada; Ɂehdzo Got’ȟ nę Gots’é Nákedı (Sahtu Renewable Resources Board), Tulita, Northwest Territories, Canada; Government of the Northwest Territories, Yellowknife, Northwest Territories, Canada; Fish and Wildlife Stewardship, Government of Alberta, Grand Prairie, Alberta, Canada; Fish and Wildlife Stewardship, Alberta Environment and Protected Areas, Edmonton, Alberta, Canada; Saskatchewan Government, Regina, Saskatchewan, Canada; Government of Manitoba, Winnipeg, Manitoba, Canada

**Author notes:** **CORRESPONDING AUTHOR**, Cory Fournier, 1600 W Bank Dr. Peterborough, ON. Canada. K9L 0G2.

**Keywords:** caribou, conservation genetics, continuously distributed species, genetic connectivity, genetic diversity, genetic networks, population structure, *Rangifer tarandus*

## Abstract

The conservation of genetic diversity and connectivity is essential for the long-term persistence and adaptive ability of a species. Recent calls have been made for the inclusion of genetic diversity and differentiation measures in the assessment, management, and conservation of species. However, the literature often lacks direction on how to do so for species with continuous distributions or no distinct breaks in genetic connectivity. There are many considerations to overcome when investigating genetic diversity and connectivity of such species. We combine multiple genetic network methodologies with more traditional population genetic analyses within a single framework to address the challenges of investigating population structure and quantifying variation in genetic diversity and connectivity of wide-ranging species with continuous distributions. We demonstrate the efficacy and applicability of our framework through a study on woodland caribou (*Rangifer tarandus*) occupying the boreal forest of Canada; a species of significant conservation concern. The dataset consisted of 4911 unique individuals genotyped at 9 microsatellite loci, which were subsequently partitioned into 103 spatial nodes to create a population-based genetic network. The Walktrap community detection algorithm was used to detect hierarchical population genetic structure within the study area and node-based network metrics such as mean inverse edge weight and clustering coefficient were used to quantify the variation in genetic connectivity across the range. Lastly, genetic diversity was assessed by calculating allelic richness and heterozygosity of the nodes making up the network. The community detection analysis identified two communities at the coarsest scale to nine communities at the optimal partition. A strong pattern of Isolation by Distance (IBD) was found across the range at multiple scales. Furthermore, signs of genetic erosion along the study area’s southern boundaries were depicted by nodes presenting low genetic diversity and low centrality values. These results are important to the species status assessments in providing previously unavailable information on connectivity and diversity within and beyond the current local population units used in management. Our approach to quantify the patterns and extent of connectivity across the boreal range is comprehensive and could easily be adapted to other species. The results are robust and provide a solid foundation for the continued monitoring and recovery of the species.

## 1 INTRODUCTION

Maintaining connectivity and gene flow at multiple spatial scales reduces the effects of demographic and environmental stochasticity and conserves genetic diversity which can be a proxy of adaptive potential (Leimu et al. 2010; Bijlsma and Loeschcke 2012; Hoban et al. 2022; Thompson et al. 2023). Habitat fragmentation, range retraction, and the isolation of animal populations often lead to increased inbreeding and genetic drift, resulting in reduced genetic diversity, inbreeding depression, and genetic erosion (Bouzat 2010; Neaves et al. 2015; Mimura et al. 2017; Thompson et al. 2019; Kardos et al. 2023). Quantifying and monitoring the spatial variation of genetic diversity and connectivity is thus critical to species management and conservation, particularly in the face of increasing anthropogenic disturbance and climate change.

The advancement of molecular techniques has facilitated an increased interest in genetic studies in support of wildlife conservation and management (DeSalle and Amato 2004; O’Brien et al. 2022; Hoban et al. 2022; Thompson et al. 2023; Tkach and Watson 2023). For the assessment and monitoring of wide-ranging and continuously distributed species however, several challenges have been identified including: i) the difficulty of determining genetic structure in the presence of Isolation by Distance (IBD) (Perez et al. 2018; Scribner et al. 2005), ii) the need for *a priori* partitioning of demes or populations to calculate genetic diversity and differentiation measures (Dyer and Nason 2004), and iii) the complexity of interpreting genetic diversity and differentiation measures in systems with multiple phylogenetic backgrounds, introgression, and divergence (Hewitt 2004; Campbell et al. 2015; Taylor et al. 2021).

Traditional approaches used to investigate population genetic structure and connectivity are often limited to calculating a single or few summary statistics between dyads of populations, which often does not adequately capture the complexity of all interpopulation relationships (Dyer and Nason 2004; Dyer 2015). Recent developments in genetic network analysis offer an alternative approach, which is flexible, can account for genetic variation across multiple spatial and temporal scales, and can seamlessly incorporate a variety of downstream analyses (Dyer and Nason 2004; Garroway et al. 2008; Dyer 2015; Koen et al. 2016; Jones and Manseau 2022, Jones et al. 2023). Population-based genetic networks, also referred to as population graphs (Dyer and Nason 2004), inform genetic covariance structure among populations or nodes of individuals. They describe the genetic variation among all populations or nodes of individuals simultaneously, rather than in a pairwise fashion, making it possible to capture the evolutionary consequences of various historical and contemporary processes. Researchers can use genetic networks to test hypotheses relating to genetic differentiation, IBD, population assignment, and genetic connectivity under a single framework (Dyer and Nason 2004). Genetic networks also provide flexibility for various downstream analyses such as running community detection algorithms and calculating node-based metrics, allowing researchers to simultaneously estimate population genetic structure and quantify the spatial variation of gene flow and connectivity (Greenbaum et al. 2016; Cross et al. 2018; Greenbaum et al. 2019; Bouchard et al. 2022).

Community detection algorithms, such as the Walktrap algorithm (Pons and Latapy 2005), partition networks into communities of nodes that have stronger and more intra-node edges than inter-node edges. Since edges in genetic networks represent genetic connectivity, the community partitions determined from the community detection algorithms can indicate signals of genetic structure (Greenbaum et al. 2019; Greenbaum et al. 2016; Savary et al. 2021a). Additionally, node-based network metrics can be used to quantify the variation in connectivity between nodes that make up a network (Jones and Manseau 2022). Metrics such as closeness centrality, mean inverse edge weight, and clustering coefficient can be used to describe how a node is connected within a network and therefore quantify the genetic or functional connectivity of the nodes within a genetic network (Cross et al. 2018; Bouchard et al. 2022). The flexibility and wide scope of genetic networks make them ideal for studying genetic diversity and connectivity of wide-ranging species.

In this study, we address the above challenges of investigating population genetic structure and quantifying variation in genetic diversity and connectivity in continuously distributed species. We achieve this by combining multiple genetic network methodologies with more traditional population genetic analyses within a single framework (Figure 1). We apply this framework to an extensive dataset (4911 unique genotypes) of two woodland caribou (*Rangifer tarandus caribou*) ecotypes occupying the Canadian boreal forest (i.e., boreal and eastern migratory caribou; COSEWIC 2011). Caribou occupying this region present a largely continuous distribution and different phylogenetic origins shaped by past glacial cycles and patterns of introgression and divergence (Klütsch et al. 2016; Polfus et al. 2017; Taylor et al. 2021; Solmundson et al. 2023). Wide-ranging species with continuous distributions often have no obvious breaks in population genetic structure to use to *a priori* partition the dataset into nodes (Perez et al. 2018). Our framework addresses this by first clustering samples within the study area at a set geographic scale (i.e.,100km) to create spatial nodes used to construct population-based genetic networks (i.e., population graphs; Dyer and Nason 2004). Building upon the “population graph” method pioneered by Dyer and Nason, we incorporate further downstream analyses on the network: community detection algorithms are used to explore genetic structure and differentiation, and node-based network metrics are calculated to simultaneously quantify variations in genetic connectivity and gene flow. Additionally, more traditional population genetic methods are applied such as testing for IBD, calculating genetic differentiation, and measures of genetic diversity; all of which are interpreted more meaningfully by incorporating community detection and node-based metric results.

**Figure 1.**
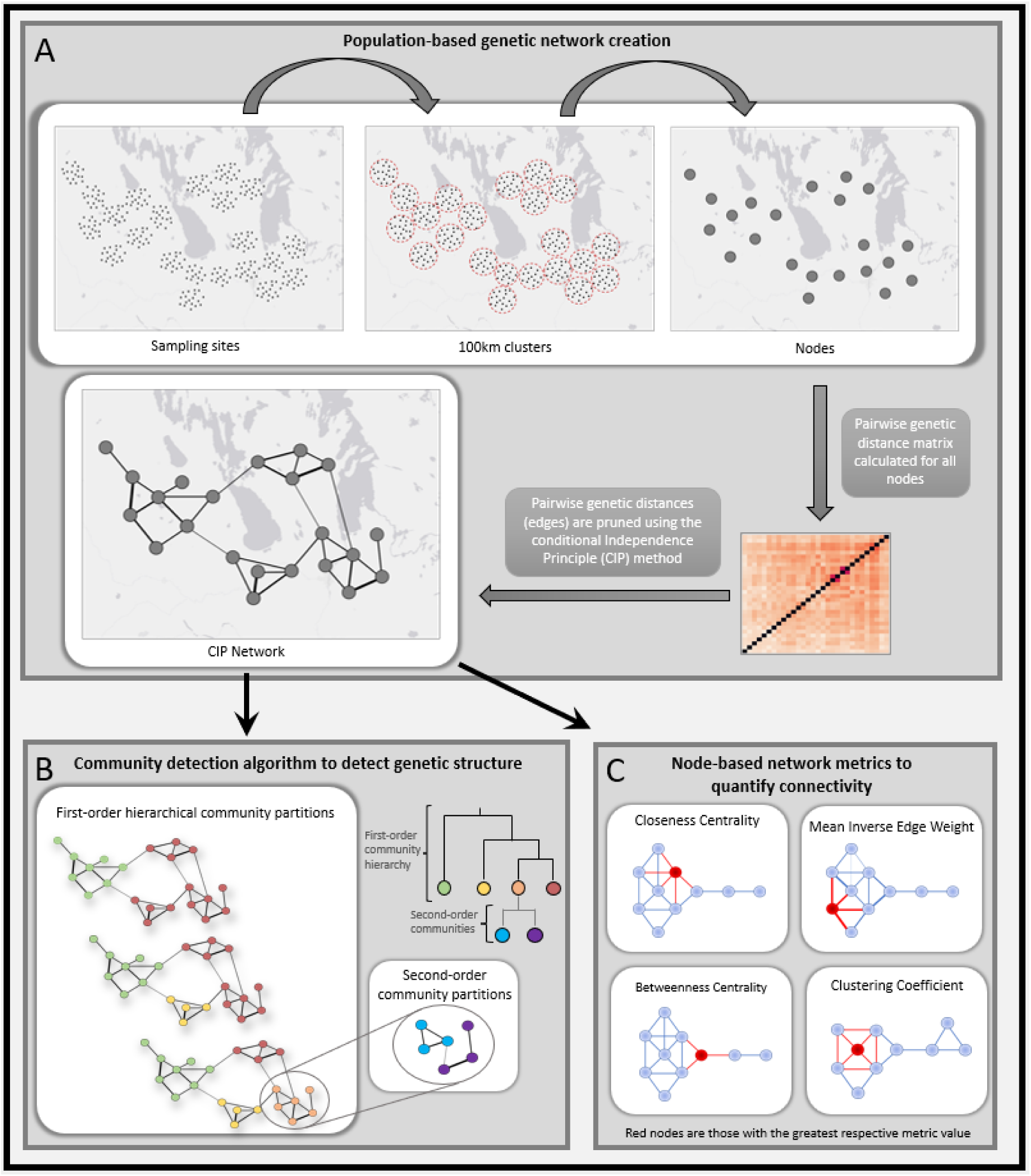
Schematic of creating the genetic network pruned using the Conditional Independence Principle (CIP) method (A) and the subsequent downstream analyses: community detection to find genetic structure (B) and node-based network metrics to quantify variations in connectivity (C). All networks in schematic are hypothetical.

Boreal caribou were federally listed as *Threatened* in 2003 under Canada’s Species-at-Risk Act (SARA) (Environment and Climate Change Canada 2020). As a central component of the recovery strategy (Environment and Climate Change Canada 2020), boreal caribou are delineated as 51 distinct local population units (LPUs) informed through telemetry, observational data, and in some cases jurisdictional boundaries (Environment Canada 2011). There is evidence that these ranges do not reflect true population boundaries as several studies have documented strong IBD and limited genetic differentiation across ranges (Ball et al. 2010; Galpern et al. 2012, Priadka et al. 2019). There is evidence of northward range retraction in Ontario resulting in increased isolation of southern populations (Schaefer 2003) and genetic erosion along the southern periphery of the range (Thompson et al. 2019). In this analysis spanning 35 of the 51 boreal caribou LPUs and the adjacent eastern migratory ecotype range, we predict nonalignment of the clustering results with the current LPUs, along with a weaker pattern of IBD within each cluster. We also predict lower genetic diversity and connectivity along the southern range margin as captured by weakly connected or disconnected nodes, nodes presenting low centrality values, and low genetic diversity. Finally, we predict to see the influence of the known evolutionary history on the variation of genetic diversity. Our analytical framework addresses the main challenges identified when dealing with the assessment and monitoring of wide-ranging and continuously distributed species. The results are critical to status assessments and in providing a robust and solid foundation for the continued monitoring and recovery of the species.

## 2 MATERIALS AND METHODS

### 2.1 Sample Collection and Genotyping

This study used a dataset of 4911 individual caribou genotypes. The study area spanned the Boreal Woodland Caribou Designatable Unit range (COSEWIC 2011) from the Northwest Territories to northeastern Ontario and included part of the Eastern Migratory Designatable Unit range (COSEWIC 2011) in Ontario and Manitoba. Our study area covered approximately 1.8 million km^2^ (see Figure 2 for a map of the study area and sampling locations). We opted to not include data from the discontinuous Little Smoky LPU in Alberta as this population clustered with mountain caribou ecotypes in previous studies (Cavedon et al. 2022; unpublished results). Genotypes were obtained from fecal samples collected as part of large-scale sampling programs over 17 years (2004-2021; www.EcogenomicsCanada.ca). All samples were amplified and analyzed at 9 microsatellite loci [BM848, BM888, MAP2C, RT24, RT30, RT5, RT6, RT7, RT9 (Bishop et al. 1994; Wilson et al. 1997; Mcloughlin et al. 2004; Cronin et al. 2005)] and animal sex was determined by utilizing Zfy/Zfx primers according to protocols outlined in Ball et al. (2007) and Klütsch et al. (2016).

**Figure 2.**
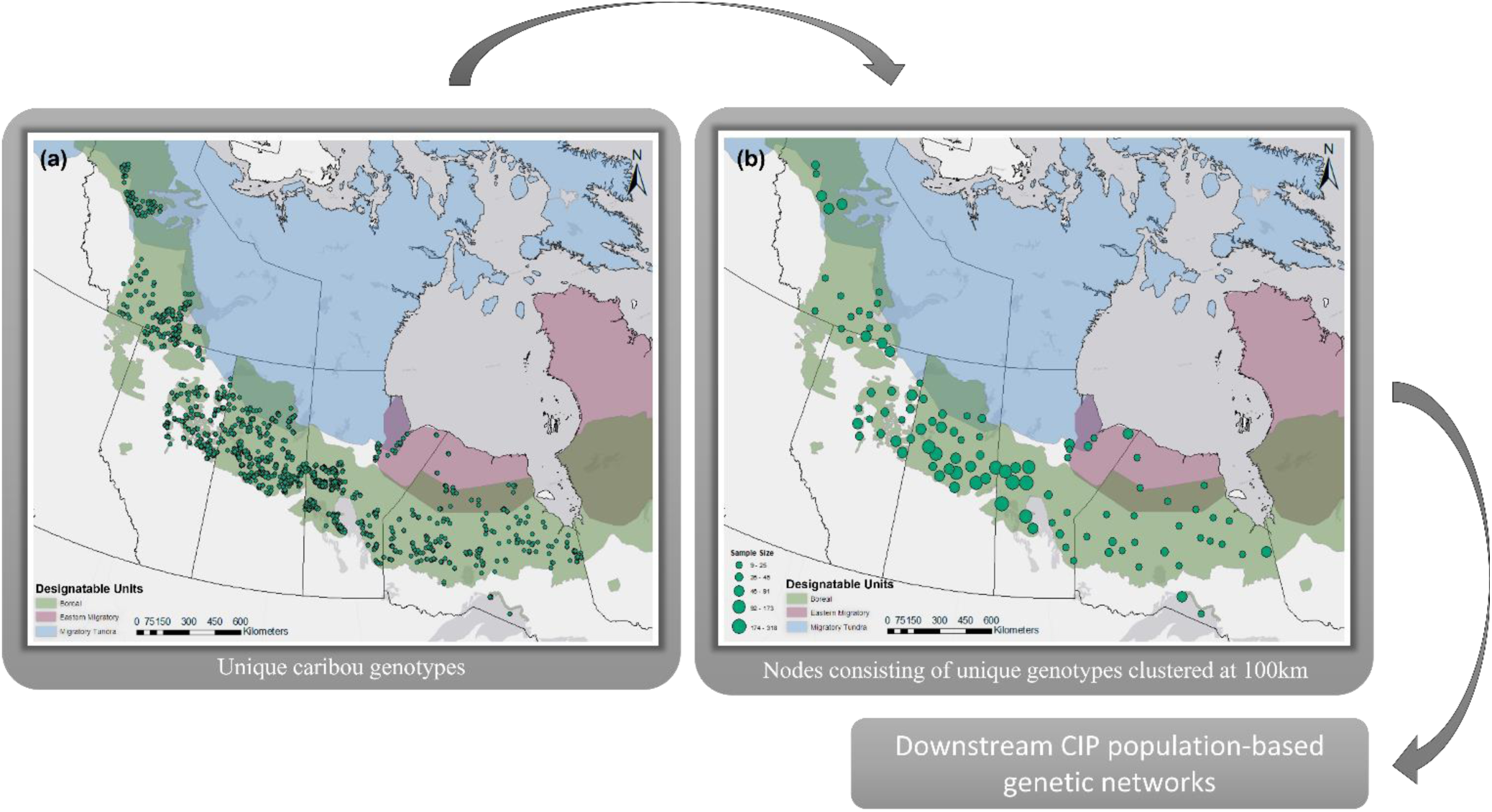
Sample locations of all 4911 unique caribou genotypes across the study area (A). Samples clustered at a 100km spatial scale to create nodes for downstream network analyses with node size representing sample size (B).

### 2.2 Genetic Network Creation

For the population-based genetic networks, we first created nodes by clustering all samples within a 100km radius (Figure 2). Acknowledging the trade-off between the number of individuals within each node and the number of nodes included in the population-based network, we opted to use a 100km scale which allowed us to maximize the number of nodes with a sample size greater than ten individuals. We achieved this by hierarchically clustering the samples based on a geographic distance matrix and then defining the clusters based on a tree height cut-off value of 100km using the *hclust* and *cutree* functions within the R Core *stats* package (R Core Team 2022). We determined the centroid of each cluster using the *rgeos* R package (Bivand and Rundel 2021) and used it as the node location for the spatial network.

The population-based genetic network was constructed using the nodes described above and Euclidean pairwise genetic distances (based on differences in allele frequencies) as edges between nodes (Dyer and Nason 2004). The genetic network began fully saturated, having an edge between every pair of nodes. The network was then pruned using the Conditional Independence Principle (CIP) method (Dyer and Nason (2004) which sets out to prune the graph down to the smallest edge set that sufficiently describes the among-population genetic covariance structure. To accurately describe the population-based genetic network that was pruned using the CIP method we will henceforth refer to it as the CIP network. The CIP network was constructed and pruned using the R package *Graph4lg* (Savary et al. 2021b) which implements functions from the R packages *Popgraph* (Dyer and Nason 2004) and *igraph* (Csardi and Nepusz 2006) to construct and analyze genetic networks (Figure 1).

### 2.3 Genetic Population Structure: Network Community Detection

To assess genetic population structure across the study area, we implemented the Walktrap community detection algorithm (Pons and Latapy 2005) on the CIP network. The Walktrap algorithm detects communities of nodes with shorter intra- and longer inter-node distances based on random walks. Since edges represent genetic distances between nodes, community partitions correspond to genetic structure within the study area represented by the network. The community detection algorithm partitions communities hierarchically, starting at two communities and increasing the number of partitions until the optimal number of communities is found. We refer to the communities identified at the optimal partition as “first-order communities.” To quantify genetic differentiation among the clusters, we calculated pairwise F_ST_ values between all first-order communities using GenAlEx 6.5 (Peakall and Smouse 2012). We additionally tested for further signatures of fine-scale genetic structure below the first-order communities. For each first-order community, we re-created CIP networks and reran the community detection algorithm using the same methods described above which further partitioned the network into “second-order communities”.

### 2.4 Quantifying variation in genetic connectivity: node-based network metrics

CIP networks represent the genetic covariance structure and describe gene flow among the nodes that make up the network. To quantify the differences in connectivity between the nodes within the network we calculated a series of node-based network metrics. Network metrics describe how a node is connected within a network and therefore quantify the genetic or functional connectivity of the nodes within a CIP network. We calculated four node-based network metrics: closeness centrality, betweenness centrality, mean inverse edge weight, and clustering coefficient. Closeness centrality is the reciprocal of the sum of the shortest path between the given node and all other nodes (Cross et al. 2018). We calculated closeness centrality to identify nodes that are well connected to *all* other nodes in the network. Betweenness centrality is defined by the number of shortest paths between all nodes that travel through the given node and all other nodes (Koen et al. 2016; Garroway et al. 2008; Cross et al. 2018). We calculated betweenness to reveal nodes that were acting as bridges or bottlenecks between otherwise disconnected components of the network therefore identifying nodes that were important for maintaining network connectivity. Mean inverse edge weight (MIW) is the mean of the inverse weights of all edges connected to the given node (Koen et al. 2016). We calculated MIW to identify nodes that had strong immediate connections to their neighboring nodes, indicating nodes having a high degree of gene flow. Lastly, clustering coefficient is defined by the probability that two nodes connected to a neighbouring node are also connected (Bouchard et al. 2022). We calculated clustering coefficient to identify nodes that may have been acting as an anchor of interconnected groups of nodes, therefore indicating nodes that were in regions of the network that were displaying stronger signals of structure.

We calculated these four metrics for each node in the CIP network. We created histograms to visualize the distribution of each metric value. For each of the four metrics, we visualized the CIP networks both spatially and non-spatially, with node sizes representing the node’s respective metric value, and identified nodes within the top ten percent of each metric.

### 2.5 Isolation by Distance (IBD)

To test for signatures of IBD, we tested for a significant correlation of the genetic and geographic distances between all nodes (Van Strien et al. 2015). We created a genetic distance matrix (pairwise F_ST_) and geographic distance matrix for all nodes using GenAlEx 6.5 (Peakall and Smouse 2012). We ran a Mantel test with 1000 permutations in R to test the correlation and associated significance between the two matrices. In addition to the Mantel test, we also calculated mantel correlograms with 10,000 permutations using the *ngram* function in the R package *ecodist* (Goslee and Urban 2007) to test for a significant correlation between genetic and geographic distance at various geographic distance intervals (intervals of 100km). To test for patterns of IBD at multiple spatial scales, using the same methods we additionally tested for IBD in the eastern and western communities of the network (coarsest hierarchical community detection partition) as well as within each of the nine first-order communities (see community detection results for more details on network partitions).

### 2.6 Genetic Diversity

Genetic diversity metrics are sensitive to sample size, a problem that rarefaction has been shown to resolve (Kalinowski 2004; Bishop et al. 2023). We therefore removed any nodes with sample sizes less than 15 and randomly subsampled larger nodes down to 15 to have consistency in sample size. We then calculated the number of alleles (Na), observed heterozygosity (Ho), and expected heterozygosity (He) for each node within GenAlEx 6.5 (Peakall and Smouse 2012). We created interpolated raster surfaces for each metric using the inverse distance weighted (IDW) technique within ESRI’s ArcMap 10.8 GIS software. We created the interpolated raster surfaces for the entire study area as well as for the eastern and western portions of the study area separately to account for the large-scale variation in genetic diversity possibly driven by the different phylogenomic origins of boreal caribou (see community detection and genetic diversity results sections for details). In addition to Na, He, and Ho, we also calculated additional population genetic summary statistics including the number of effective alleles (Ne), Shannon’s Information Index (I), Unbiased Expected Heterozygosity (uHe), and the Fixation Index (F) for all nodes to provide further information on intraspecific variation of genetic diversity.

## 3 RESULTS

### 3.1 Genetic Networks

The network, created with Euclidean genetic distances as edges between the nodes and pruned with the CIP method, had 320 edges connecting the 103 nodes. The network had no disconnected components, with each node having at minimum one edge connecting it to a neighbouring node (Figure 3).

**Figure 3.**
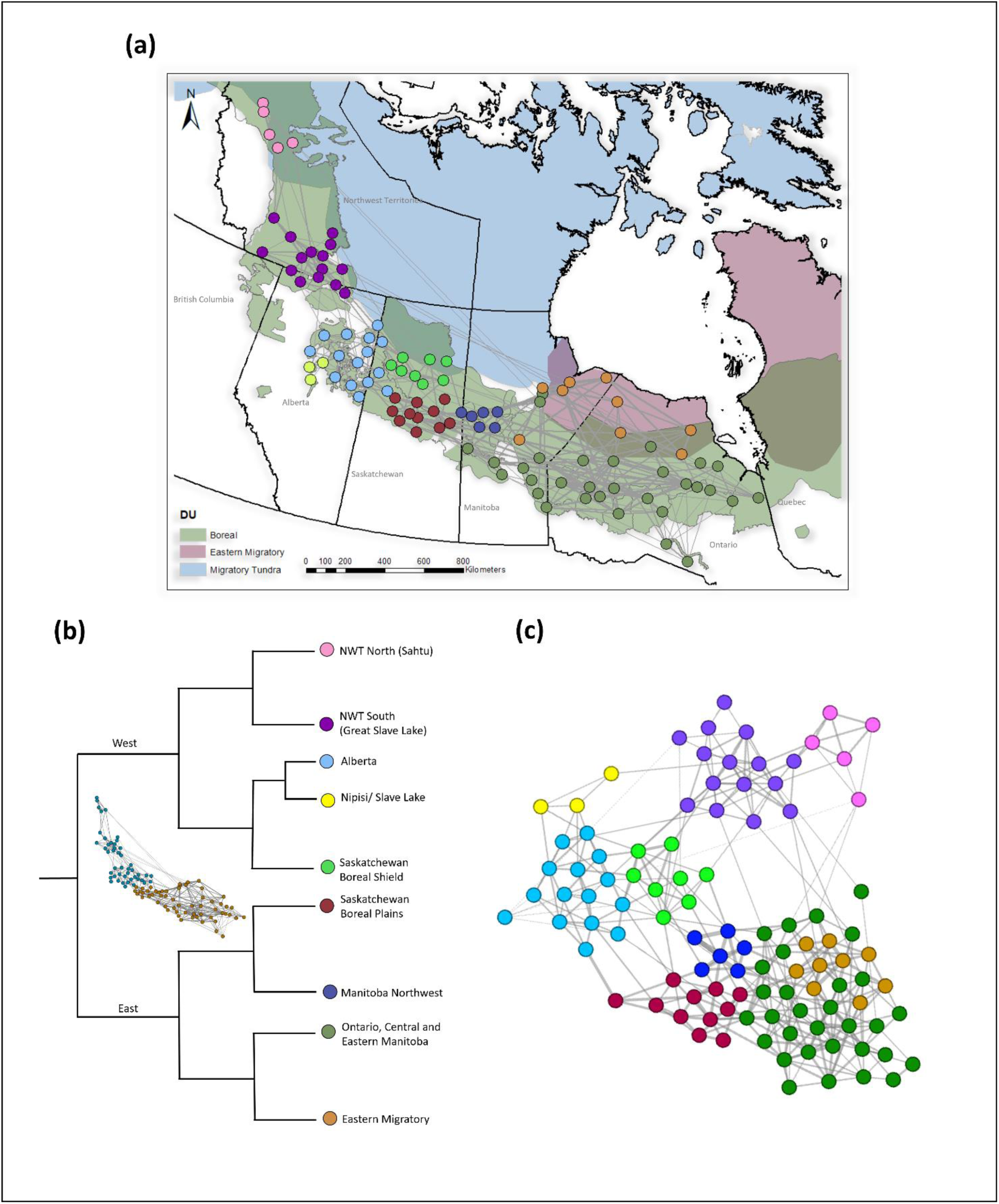
Genetic network pruned using the conditional independence priciple method (CIP Network) mapped spatially (A). Dendrogram representing heirarchical community detection of the CIP Network (B). CIP Network represented non-spatially (C). Node colours represent first-order community assignment at the optimal partition of the Walktrap community detection algorithm.

### 3.2 Community Detection

The Walktrap community detection algorithm detected nine communities at the optimal partition that were hierarchically clustered. The clustering hierarchy is represented by the dendrogram in Figure 3B. At the first partition, the Walktrap community detection algorithm partitioned the network into two large communities, comprising the eastern and western portions of the study area respectively. The spatial CIP network (Figure 3) showed spatial variation in edge density, with the eastern portion of the network having a higher edge-to-node ratio than the western portion of the network.

The western portion further separated into five first-order communities and the eastern portion separated into four first-order communities at the optimal partition (Figure 3). There were two communities in the Northwest Territories, one in the north (5 nodes; hereafter referred to as NWT North) and one in the south (15 nodes; hereafter referred to as NWT South). There was a large community made up of the majority of nodes in Alberta (15 nodes, hereafter referred to as Alberta), with three nodes comprised of the Nipisi and Slave Lake LPUs breaking off into their own community (3 nodes; hereafter referred to as Nipisi/ Slave Lake). Saskatchewan had two communities that aligned well with the Saskatchewan Boreal Shield LPU in the north (8 nodes: hereafter referred to as Saskatchewan Boreal Shield) and Saskatchewan Boreal Plains LPU in the south (11 nodes; hereafter referred to as Saskatchewan Boreal Plains).

In the eastern portion of the range where the network was more densely connected, there was a community in northwestern Manitoba (6 nodes; hereafter referred to as Manitoba Northwest), a large community comprised of Ontario, central and eastern Manitoba (31 nodes: hereafter referred to as Ontario, Central and Eastern Manitoba), and lastly, there was a community in northern Ontario and Manitoba that aligned with eastern migratory caribou (9 nodes; hereafter referred to as Eastern Migratory). The pairwise F_ST_ values between all first-order communities (Table 1) were relatively low for all spatially adjacent communities. F_ST_ values revealed increasing genetic differentiation between first-order communities as geographic distance increased between them. Notably, the Nipisi/ Slave Lake community displayed the highest levels of genetic differentiation.

**Table 1.** Genetic differentiation between first-order communities calculated with pairwise F_ST_.

| $F_{ST}$ | Ontario | NWT<br>South | SK<br>Plains | Alberta | SK<br>Shield | NWT<br>North | MB<br>Northwest | Eastern<br>Migratory | Nipisi/<br>Slave<br>Lake |
| --- | --- | --- | --- | --- | --- | --- | --- | --- | --- |
| Ontario | 0.000 |  |  |  |  |  |  |  |  |
| NWT<br>South | 0.049 | 0.000 |  |  |  |  |  |  |  |
| SK Plains | 0.011 | 0.034 | 0.000 |  |  |  |  |  |  |
| Alberta | 0.031 | 0.022 | 0.015 | 0.000 |  |  |  |  |  |
| SK Shield | 0.021 | 0.029 | 0.007 | 0.013 | 0.000 |  |  |  |  |
| NWT<br>North | 0.054 | 0.009 | 0.037 | 0.029 | 0.031 | 0.000 |  |  |  |
| MB<br>Northwest | 0.009 | 0.036 | 0.008 | 0.025 | 0.013 | 0.040 | 0.000 |  |  |
| Eastern<br>Migratory | 0.009 | 0.050 | 0.012 | 0.033 | 0.018 | 0.052 | 0.009 | 0.000 |  |
| Nipisi/<br>Slave Lake | 0.057 | 0.037 | 0.048 | 0.027 | 0.045 | 0.047 | 0.052 | 0.061 | 0.000 |

After recreating CIP networks for each of the first-order communities, the Walktrap community detection algorithm detected second-order structure below the first-order level for six of the nine first-order communities (See Figures S1-S6 in Appendix S1).

### 3.3 Quantifying variation in genetic connectivity: node-based network metrics

Closeness centrality values were greater in the eastern portion of the network than in the western portion (Figure 4A/B). Nine of the ten nodes within the top ten percent of closeness centrality values were found within the eastern portion of the network. Mean Inverse Edge Weight (MIW) values were again greater within the eastern portion of the range compared to the western portion (Figure 4C/D). All nodes within the top ten percent of MIW values were found within the eastern portion of the range. Nodes with high betweenness centrality values were found at the interface of the east and west portions of the network in Saskatchewan as well as at the interface between the Northwest Territories and Alberta (Figure 4E/F). Lastly, nodes with the highest clustering coefficients were primarily located in the western portion of the range (Figure 4G/H).

**Figure 4.**
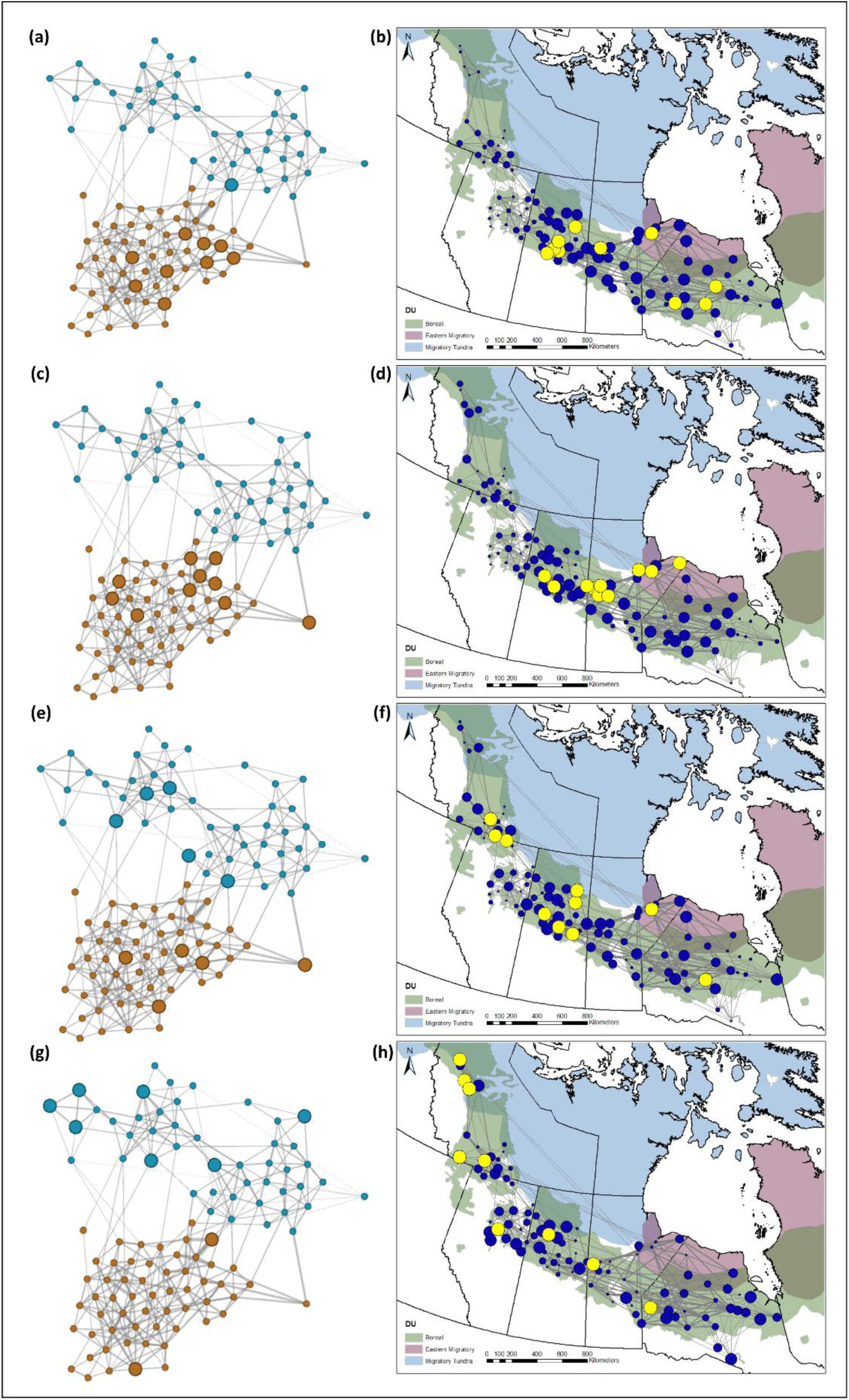
Genetic networks pruned with the conditional independence priciple method representing variation in connectivity through node-based metric values: Closeness Centrality (A/B), Mean Inverse Edge Weight (C/D), Betweenness Centraltiy (E/F) and Clustering Coefficient (G/H). Aspatial networks (A/C/E/G) have the western portion of the network represented by blue nodes, the eastern portion of the network represented by brown nodes, and the top ten percent of each metric identified with larger nodes. The spatial networks (B/D/E/H) depict the respective metric values with node size and highlight the top ten percent of each metric in yellow.

### 3.4 Isolation by Distance

A significant pattern of isolation by distance (IBD) was found across the entire study area based on a Mantel test between the genetic and geographic distance matrixes of all nodes (*r*=0.62, *p*=0.001) (Figure 5A). The Mantel correlogram detected a significant correlation between genetic and geographic distances at 100km distance intervals up to 900km (Figure 5B). When we tested for IBD within the eastern and western communities separately, significant correlations (IBD) were detected for each community (east: *r*=0.32, *p*=0.001; west: *r*=0.35, *p*=0.001). In the west, the correlation between genetic and geographic distances was much stronger at shorter distances (up to 300km), after which the correlation rapidly plateaued (Figure 5C/D). In the east, genetic distances increased more monotonically with increasing geographic distances and remained significant up to 500km (Figure 5E/F). At the finest scale, we tested for IBD within each first-order community and found significant patterns of IBD in five of the nine communities (Table 2). First-order communities in the east had a generally stronger signal of IBD, apart from the Eastern Migratory community. In the west, the first-order communities displayed either no significant IBD or very weak correlations.

**Figure 5.**
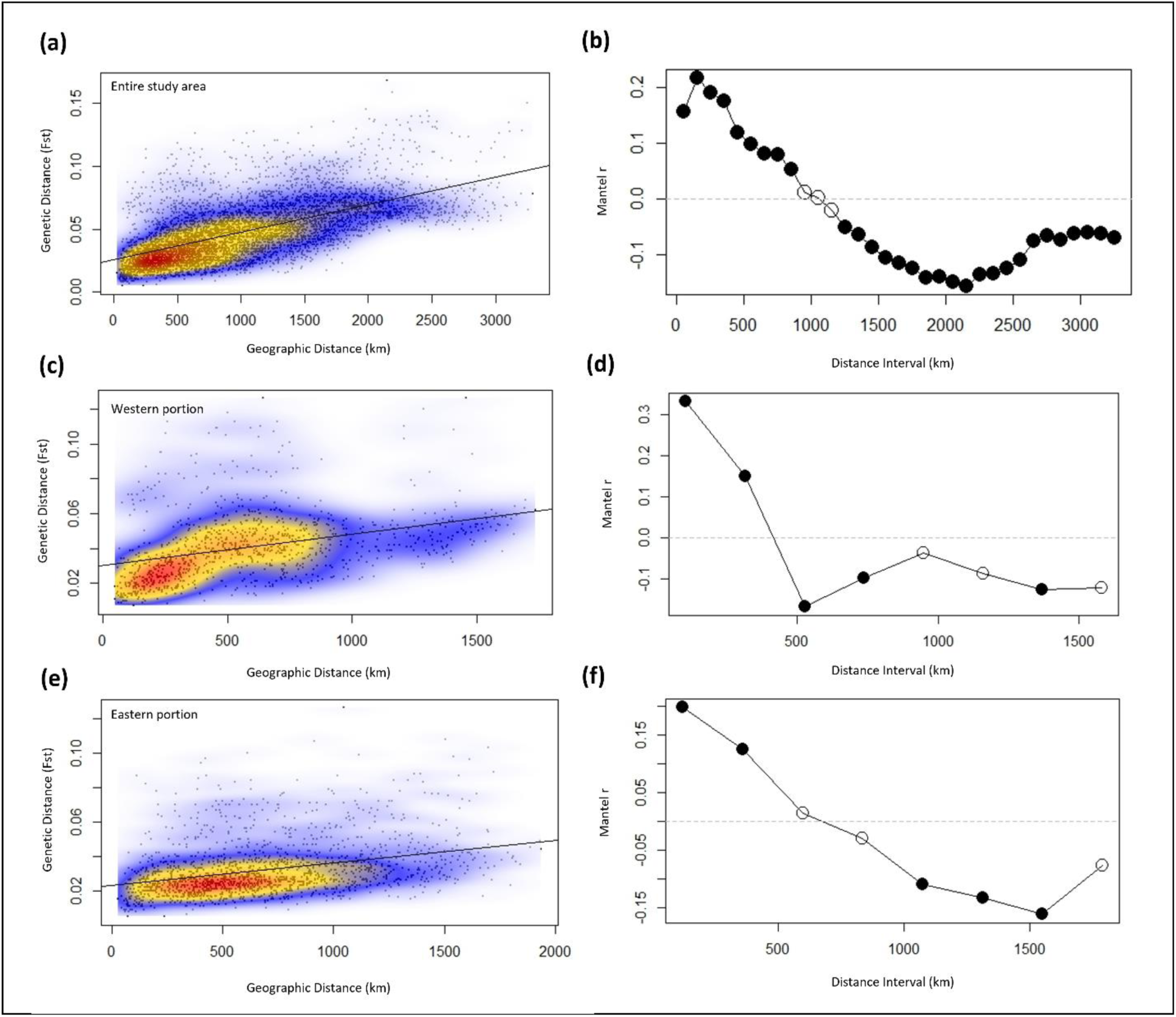
Isolation by distance (IBD) scatter plots (A/C/E) depicting the results of a Mantel test on the genetic and geographic distance matrices between all 103 nodes used to create the genetic networks. Colours represent the relative density of points on the plot: red high density, yellow medium density, and blue low density. Isolation by distance (IBD) Mantel Correlograms (B/D/F) depicting the correlation between the genetic and geographic distance matrices at various geographic distance intervals. Solid circles represent significant correlation at the respective distance interval. Mantel test and mantel correlograms were computed for the entire study area (A/B), the western portion of the study area (C/D), and the eastern portion of the study area (E/F).

**Table 2.** Mantel test results for the correlation between geographic and genetic distance matrices (isolation by distance) of the nodes within each first-order community.

| **First-order community** | **r** | **p-value** |
| --- | --- | --- |
| Ontario | 0.258* | 0.004 |
| NWT South/ Great Slave Lake | 0.296* | 0.049 |
| SK2 | 0.362* | 0.018 |
| Alberta | -0.022 | 0.495 |
| SK1 | 0.499* | 0.004 |
| NWT North/ Sahtu | -0.043 | 0.548 |
| Manitoba North | 0.824* | 0.007 |
| Eastern Migratory | 0.068 | 0.361 |
| Nipisi/ Slave Lake | 0.936 | 0.33 |

\* represents significant correlation ( $\alpha = 0.05$ )

### 3.5 Genetic Diversity

The genetic diversity parameters (Na, He, and Ho) calculated for the entire study area displayed an east-to-west pattern of spatial variation (Figure 6A/B) with the east having generally lower genetic diversity compared to the west. This east-to-west pattern of genetic diversity variation reflected the phylogenomic signature that was also captured by the two large network communities. Summary statistics (Na, He, Ho, Ne, uHe, I, and F) for each node can be found in Appendix S1 (Table S2).

**Figure 6.**
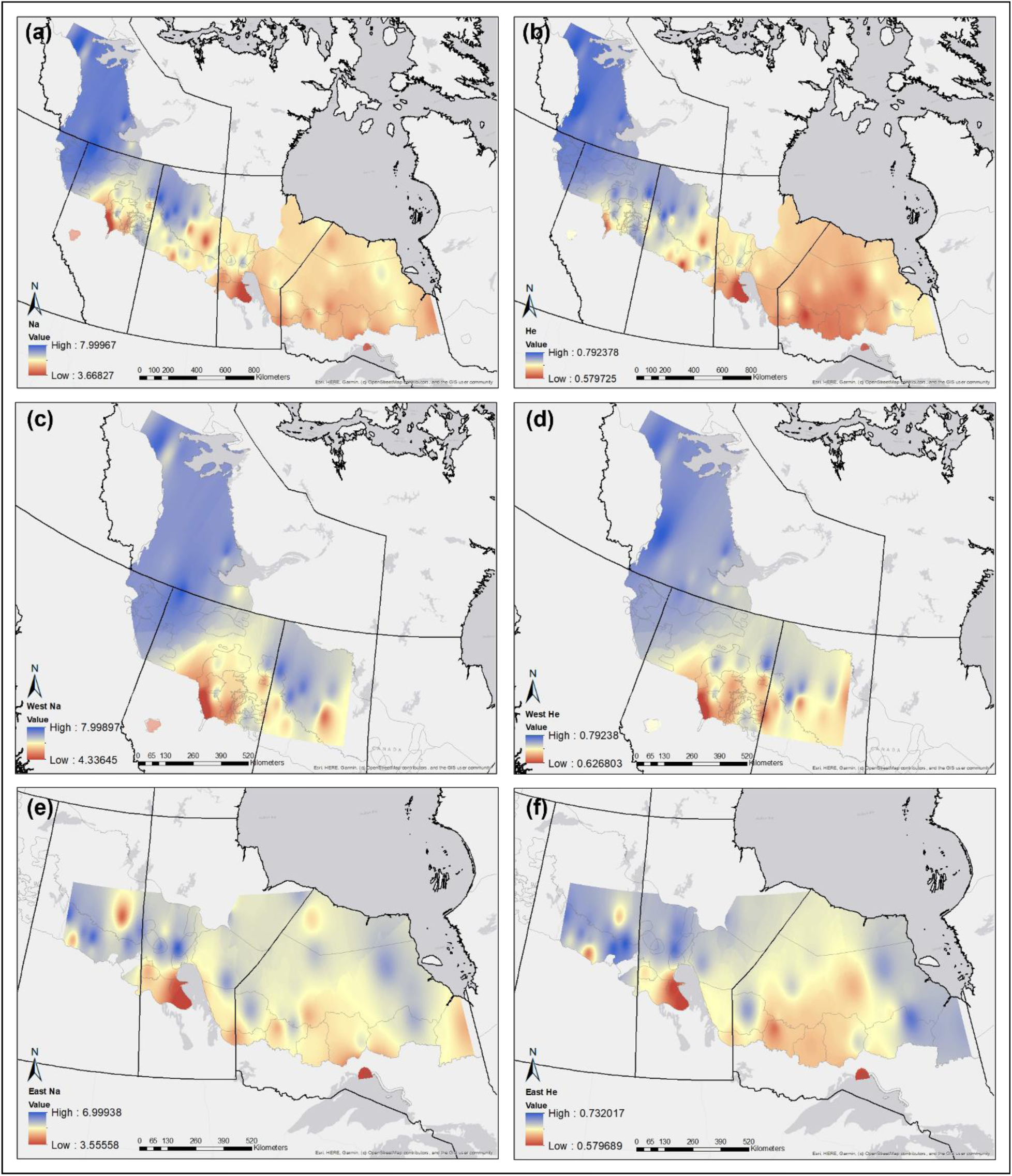
Maps of interpolated raster surfaces of two different measures of genetic diversity: Number of Alleles (Na; A/C/E) and Expected Heterozygosity (He; B/D/F). Interpolated raster surfaces representing variation in Na and He are first depicted across the entire study area (A/B). Interpolated raster surfaces representing variation in Na and He are then depicted for the western portion of the range (C/D) and eastern portion of the range (E/F) separately. Green high genetic diversity, yellow medium genetic diversity, and red low genetic diversity.

Once the phylogenomic patterns of genetic diversity were accounted for by creating individual interpolated raster surfaces for the eastern and western portions of the range, a clear pattern of lower genetic diversity became evident in the southern periphery of the range in both the east and the west. In the west, lower values of Na and He were found along the southern edge of the range, particularly in the Nipisi and Slave Lake regions of Alberta (Figure 6C/D). In the east, lower values of Na and He were also found along the southern edge of the distribution, particularly in the north Interlake region of Manitoba, the Lake Superior discontinuous coastal population in Ontario, and a large southern portion of the intact boreal range in Ontario (Figure 6E/F).

## 4 DISCUSSION

We applied a framework combining multiple network-based methodologies to investigate variation in genetic diversity and connectivity of a species presenting a largely continuous distribution. Using a large boreal and eastern migratory caribou dataset, we demonstrated that the framework can overcome challenges when dealing with wide-ranging species presenting continuous distributions. To our knowledge, this is the first study to examine intraspecific variation in genetic connectivity and diversity of boreal caribou across such a large spatial extent, using such an extensive dataset. The framework first begins by clustering the samples based on a set geographic distance to create the spatial nodes for the population-based genetic networks. This is a critical step that can be used when *a priori* population structure is not known; a challenge often overlooked in literature advocating for the use of population genetic diversity and differentiation measures in the assessment, monitoring, and conservation of wild species (O’Brien et al. 2022; Hoban et al. 2022). In our caribou study, we opted for a geographic distance of 100km to cluster the samples; resulting in the 4911 samples being clustered into 103 nodes. Once the CIP network was created using the 103 nodes, the resulting network was highly connected, with no disconnected components suggesting gene flow occurring over large geographic scales. The highly connected single-component network coupled with the strong pattern of IBD found across the study area at multiple scales strengthened our assumption that caribou within the study area were largely continuously distributed, with no distinct breaks in genetic connectivity.

### 4.1 Hierarchical community detection: uncovering population genetic structure

A major component of the downstream network analysis was the community detection algorithm used to partition the CIP network into hierarchical communities. These communities infer hierarchical population genetic structure and variation in genetic connectivity at different spatial scales, likely shaped by various historical and contemporary processes. Notably, at the first hierarchical community partition, the east-to-west partition aligned well with the known phylogenomic signature of boreal caribou (Polfus et al. 2017; Taylor et al. 2021). This coarse community partition likely represents genetic structure that was shaped by historical processes such as vicariance and climactic/glacial cycles (Weksler et al. 2010; Polfus et al. 2017; Taylor et al. 2021). Although we did not empirically test if this partition was due solely to the different phylogenetic origins of caribou in this region, population graphs for other species revealed similar phylogenetic patterns (Dyer and Nason 2004). At a finer scale, the optimal community partition of the CIP network revealed nine first-order genetic communities. The community partitions at both the coarsest scale (2 communities) and optimal scale (9 communities) aligned with results from an unpublished report that used the well-known Bayesian clustering algorithm implemented in the program STRUCTURE (Pritchard et al. 2000). Although the first-order communities partially aligned with certain boreal caribou LPUs (Saskatchewan Plains and Saskatchewan Shield; Environment and Climate Change Canada 2020), they were generally larger than many of the currently delineated LPUs. The first-order communities often spanned multiple LPUs, indicating that gene flow occurred at larger scales than the currently identified boreal caribou local populations (Environment and Climate Change Canada 2020). An exception to this pattern was in the Northwest Territories, where the one large LPU representing the entire boreal caribou range in the Territory was divided into two communities.

### 4.2 Node-based metrics: quantifying intra-specific variation in genetic connectivity

The second downstream analysis within the framework calculated various node-based metrics on the CIP network to quantify variation in genetic connectivity within the study area. There was variation among node-based metric values that aligned with the first hierarchical community partition, again indicating the population genetic structure in the region was shaped to some extent by the phylogenetic origins of boreal caribou. The greater closeness centrality and mean inverse edge weight values in the east showed evidence of the eastern portion of the network having greater genetic connectivity than the west. Nodes in the eastern portion of the network displayed higher levels of overall connectivity (revealed through greater closeness centrality values) and greater direct gene flow (revealed through greater mean inverse edge weight values) compared to the western portion of the network. Furthermore, the western portion of the network showed a stronger signal of genetic structure compared to the east, indicated by nodes with greater clustering coefficients. Although there were differences in the genetic connectivity between the eastern and western communities, the two communities still had gene flow occurring between them as revealed by edges connecting multiple high betweenness nodes belonging to each of the two communities and a relatively low measure of pairwise F_ST_ (0.02). In addition to the population genetic structure of caribou in the region being shaped by their phylogenetic origins, differences in anthropogenic impacts and natural caribou landform/ habitat discontinuities (Environment and Climate Change Canada 2020) between the eastern and western portions of the study area could have also contributed to the resulting variation in genetic connectivity and structure.

### 4.3 Variation in genetic diversity: revealing genetic erosion

In addition to the patterns of population genetic structure and spatial variation of genetic connectivity found across the study area, we also used the hierarchical community detection partitions and node-based metrics to better interpret more traditional population genetic analyses. Upon calculating the commonly used measures of genetic diversity (Na, He, and Ho), we again found that the broad-scale variation in genetic diversity across the study area captured the phylogenetic patterns. The eastern portion of the range presented overall lower genetic diversity compared to the western portion of the range. The higher baseline genetic diversity in the west could be due to animals in the Northwest Territories being of Beringian-Eurasian Lineage (BEL) origin (Polfus et al. 2017), in conjunction with overall higher levels of introgression into the North American Lineages (NAL) in the west – considering that genetic diversity and historical effective population sizes in the Beringian refugia during the Pleistocene were larger (Klütsch et al. 2016; Taylor et al. 2021; Solmundson et al. 2023). By incorporating the first hierarchical community detection partition from the CIP network, we were able to control for the overall variation of genetic diversity between the eastern and western portions of the range and create interpolated raster surfaces that revealed genetic erosion occurring along the southern range periphery in both the east and west. This genetic erosion is likely caused by range retraction and habitat fragmentation due to increased anthropogenic disturbance causing breaks in caribou continuity along the southern periphery of the study area beyond natural discontinuities in caribou landforms and habitat (Schaefer 2003; Thompson et al. 2019). Without accounting for the baseline variation in genetic diversity between the eastern and western communities, the genetic erosion along the southern periphery of the range may not have been apparent and could have been eclipsed by the lower overall genetic diversity in the east.

The genetic erosion captured in the southern range boundaries, particularly in Alberta and the Lake Superior coastal population in Ontario, was further highlighted by lower node-based metric values (closeness centrality and MIW) which could again be indicative of increased fragmentation and isolation of caribou in those regions. These results align with recent genomic studies focusing on runs of homozygosity which found evidence of increased inbreeding in Ontario’s coastal populations (Solmundson et al. 2023). The reduced levels of genetic diversity (and hence the associated reduced adaptive potential) in the southern range extent has important management implications for boreal caribou, particularly since some areas showing low genetic diversity are within LPUs (i.e. Manitoba’s North Interlake and Ontario’s Coastal populations) that are considered *likely* to be self-sustaining, the lowest level of concern within the current boreal caribou recovery strategy (Environment and Climate Change Canada 2020).

## 5 CONCLUSION

In this study, we demonstrated the efficacy of combining multiple network-based methodologies into a single framework to detect hierarchical population genetic structure and variation in genetic diversity and connectivity of a species presenting a largely continuous distribution. Our framework provides a rigorous way to address recent calls to consider genetic information in the assessment, management, and conservation of wildlife species (Thompson et al. 2023; Zimmerman et al. 2023; O’Brien et al. 2022; Hoban et al. 2022; Tkach and Watson 2023) which we believe to be broadly applicable to other wide-ranging species that have a continuous distribution and/or a strong pattern of IBD.

The results of this study are critical to the current species range assessments for boreal caribou (Environment and Climate Change Canada 2020) as they provide previously unavailable information on genetic connectivity within and beyond the currently delineated LPUs. The population genetic structure revealed through the community detection algorithm indicates there is historical and contemporary genetic connectivity that spans beyond the current units of management for local populations of boreal caribou, with more constrained connectivity in western Canada. The genetic erosion and associated increased genetic differentiation uncovered along the southern edge of the distribution, where there is the highest degree of anthropogenic impacts (Environment and Climate Change Canada 2020), is concerning in relation to boreal caribou conservation and recovery. This pattern is a striking juxtaposition with the otherwise general pattern of boreal caribou genetic continuity and diversity. This spatial variation in genetic diversity and connectivity provides further evidence of the endangerment of boreal caribou in relation to ongoing increases in anthropogenic disturbance. Seeing that gene flow and genetic connectivity are critical for species to avert the effects of demographic and environmental stochasticity, and to maintain their long-term adaptive potential, it is vital that managers and policymakers incorporate measures of genetic diversity and connectivity in current and future plans and actions to conserve and recover boreal caribou across the extent of their distribution in Canada.

## ACKNOWLEDGMENTS

We would like to thank lab personnel at Trent University for the DNA extraction and analysis; Sonesinh Keobouasone at Environment and Climate Change Canada for data management; students and staff from the EcoGenomics Canada (www.EcogenomicsCanada.ca) for insightful conversations; and numerous partners and consulting companies from across Canada involved in the aerial surveys and field sampling programs.

## AUTHOR CONTRIBUTIONS

CF: conceptualization, methodology and analytical approach, lead data analysis and manuscript writing.

MM: conceptualization, funding acquisition, design and coordination of aerial surveys, methodology and analytical approach, contributed to initial draft, reviewed/edited manuscript.

BR: DNA extraction and analysis, technical laboratory support.

LA, AK, DA, TH, GP, VT, DB: Coordinated or collected samples, reviewed/ edited manuscript.

PJW: conceptualization, funding acquisition, methodology and analytical approach, reviewed/edited manuscript.

