## Appendix S1 for "Genetic network analysis uncovers spatial variation in diversity and connectivity of a species presenting a continuous distribution"

**Genetic networks uncover spatial variation in diversity and connectivity: A case study of boreal caribou**

Ecological Applications

Cory Fournier | Micheline Manseau | Bridget Redquest | Leon Andrews | Allicia Kelly | Dave Hervieux | Troy Hegel | Gigi Pittoello | Vicki Trim | Dennis Brannen | Paul Wilson

**Figure S1**. CIP Population-based networks for the NWT South (Great Slave Lake) community with node colour representing second-order community assignment.

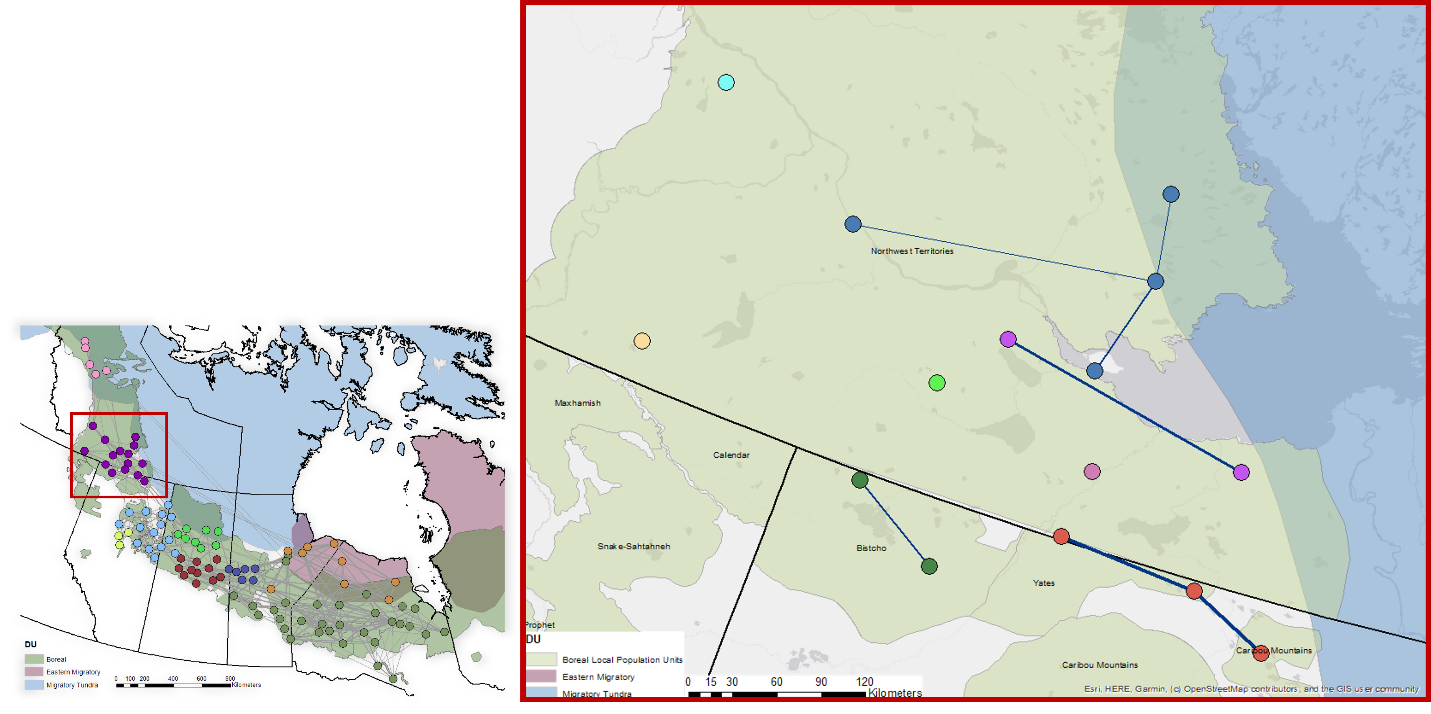

**Figure S2**. CIP Population-based networks for the Alberta community with node colour representing second-order community assignment.

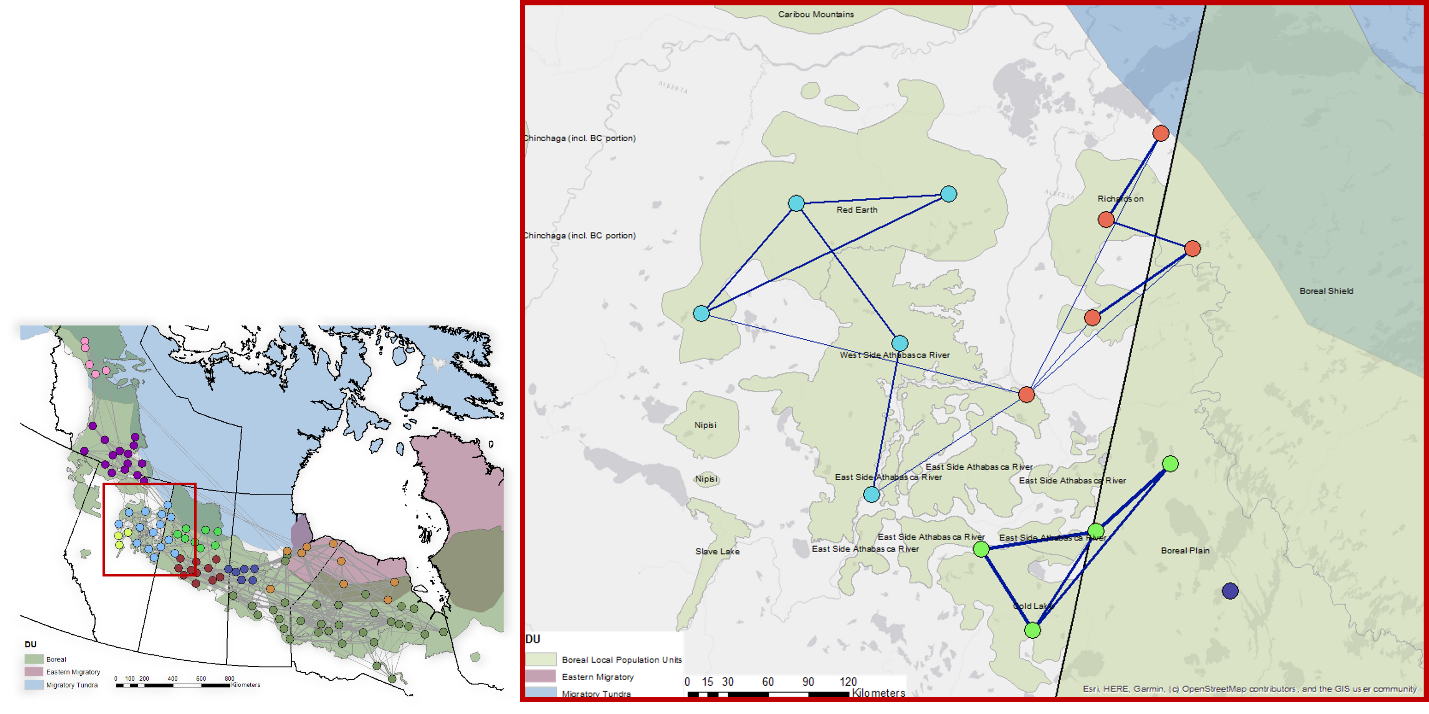

**Figure S3**. CIP Population-based networks for the Saskatchewan Boreal Shield community with node colour representing second-order community assignment.

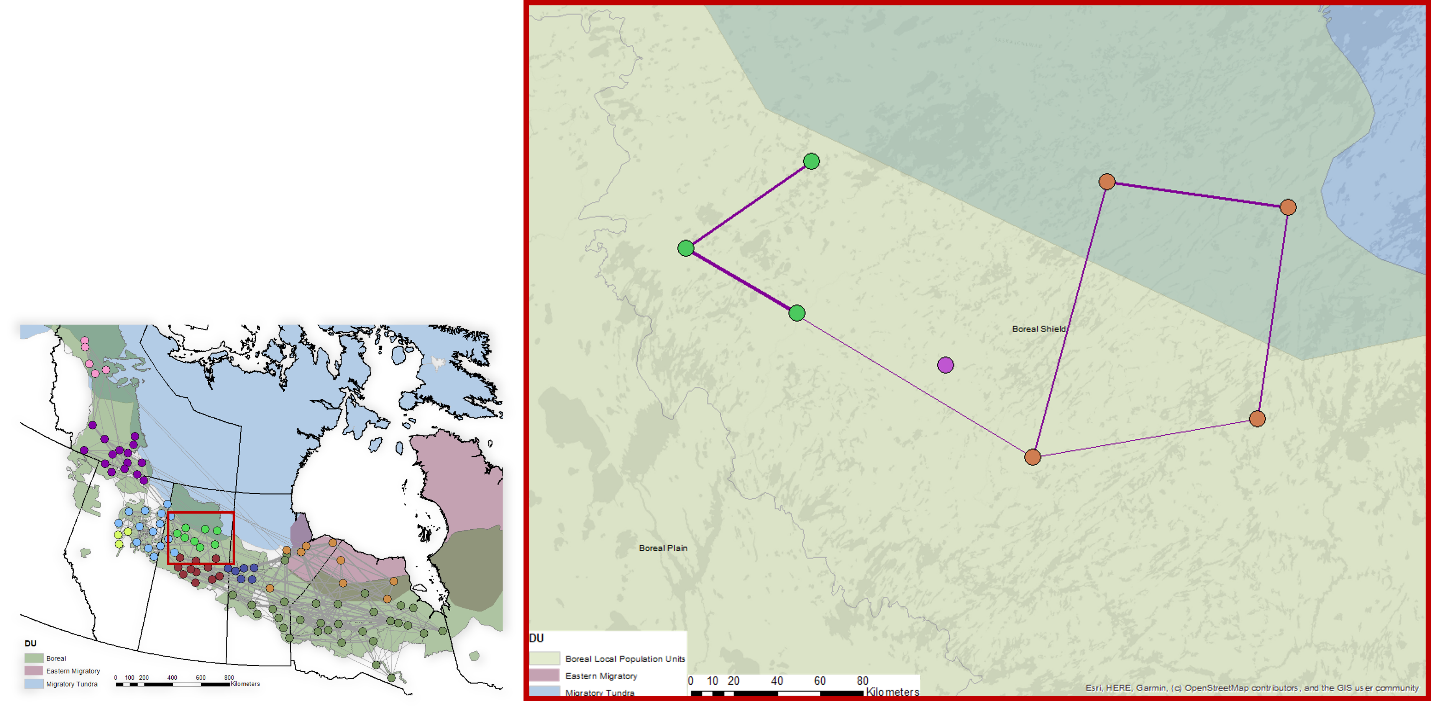

**Figure S4**. CIP Population-based networks for the Saskatchewan Boreal Plains community with node colour representing second-order community assignment.

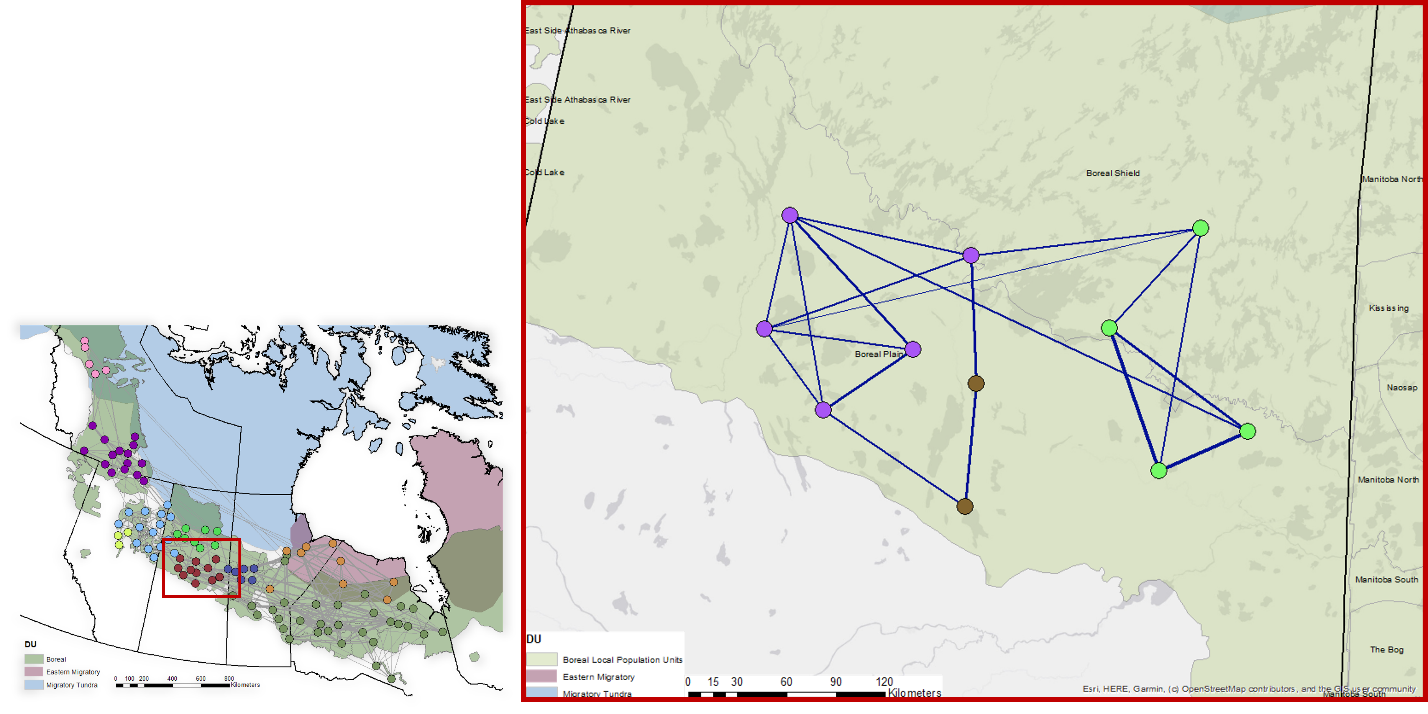

**Figure S5**. CIP Population-based networks for the Ontario, Central and Eastern Manitoba community with node colour representing second-order community assignment.

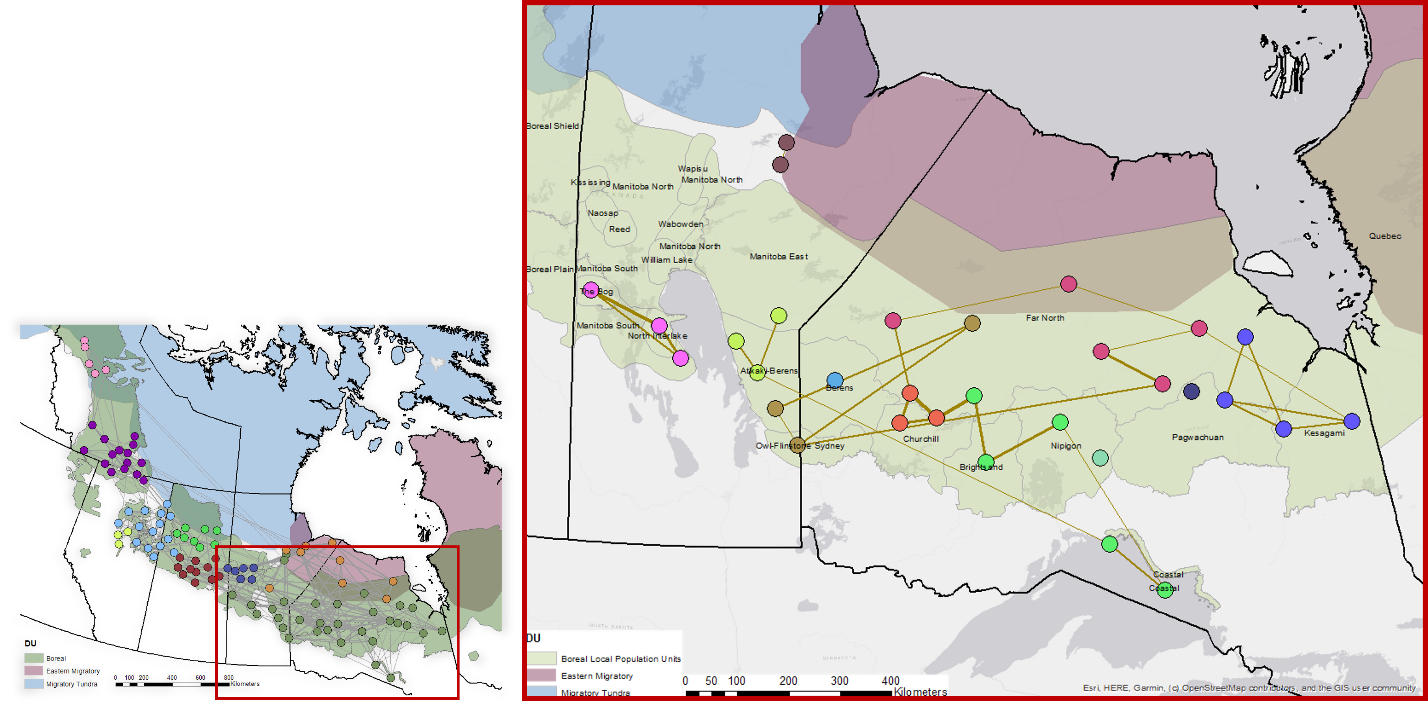

**Figure S6**. CIP Population-based networks for the Eastern Migratory community with node colour representing second-order community assignment.

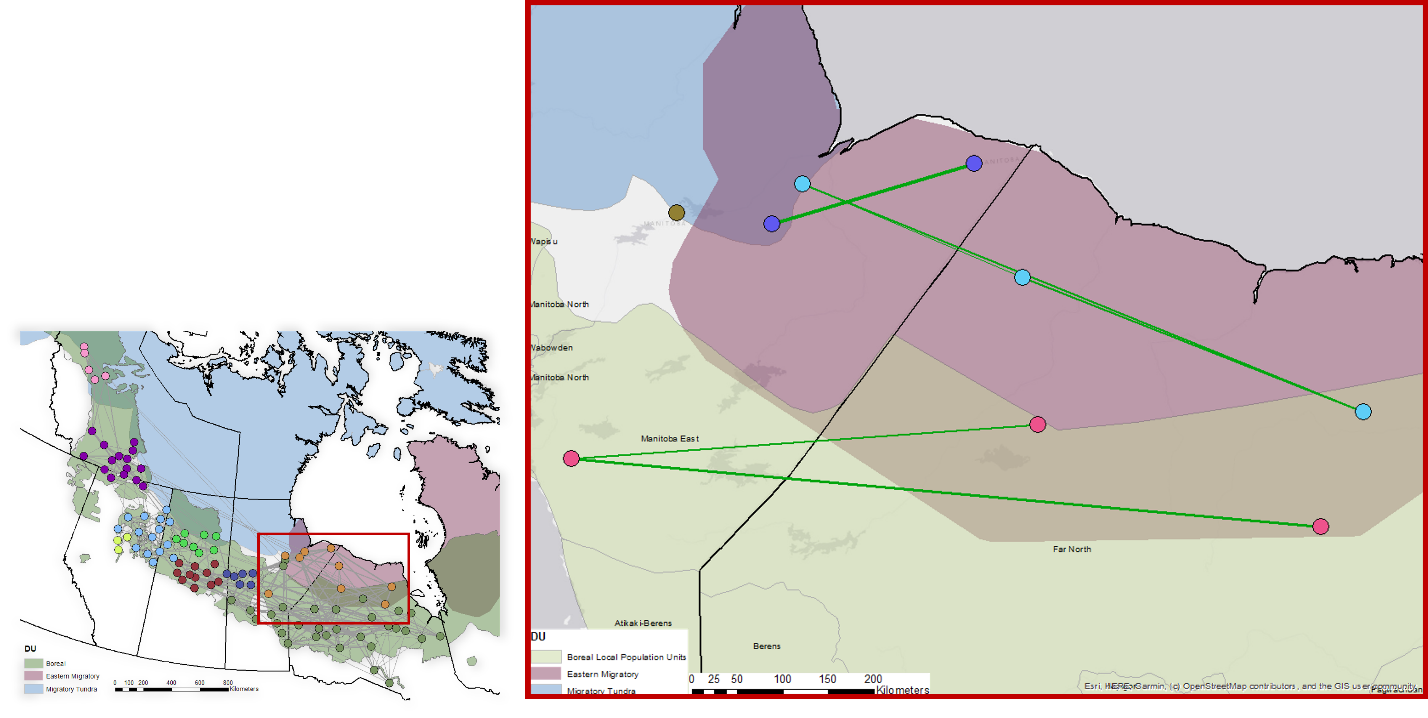

| Table S1 Node-based network metrics for all nodes in the genetic network pruned using the conditional independence principal method. Closeness centrality (close), betweenness centrality (btw), mean inverse edge weight (miw), and clustering coefficient (clust). | | | | | | | | | | | | | | | | |
| --- | --- | --- | --- | --- | --- | --- | --- | --- | --- | --- | --- | --- | --- | --- | --- | --- |
| Node | | **close** | | | **btw** | | **miw** | | | **clust** | | **Longitude** | | | **Latitude** | |
| 1 | | 0.000662 | | | 207 | | 0.225 | | | 0.107 | | -99.206734 | | | 52.84345 | |
| 2 | | 0.000649 | | | 146 | | 0.219 | | | 0.143 | | -101.27399 | | | 53.39094 | |
| 3 | | 0.000639 | | | 9 | | 0.431 | | | 0.667 | | -101.19035 | | | 54.93762 | |
| 4 | | 0.000626 | | | 122 | | 0.295 | | | 0.167 | | -99.263834 | | | 54.46166 | |
| 5 | | 0.000696 | | | 359 | | 0.262 | | | 0.071 | | -105.4728 | | | 54.59345 | |
| 6 | | 0.000686 | | | 209 | | 0.277 | | | 0.200 | | -106.12326 | | | 54.73106 | |
| 7 | | 0.000601 | | | 109 | | 0.189 | | | 0.167 | | -105.41609 | | | 53.91413 | |
| 8 | | 0.000676 | | | 238 | | 0.203 | | | 0.067 | | -106.86927 | | | 54.32258 | |
| 9 | | 0.00059 | | | 108 | | 0.193 | | | 0.200 | | -107.55323 | | | 54.7093 | |
| 10 | | 0.000648 | | | 121 | | 0.362 | | | 0.200 | | -100.49143 | | | 54.45293 | |
| 11 | | 0.000593 | | | 97 | | 0.158 | | | 0.067 | | -98.556959 | | | 52.29454 | |
| 12 | | 0.000701 | | | 255 | | 0.300 | | | 0.048 | | -100.28034 | | | 55.16079 | |
| 13 | | 0.000638 | | | 146 | | 0.260 | | | 0.067 | | -99.112956 | | | 55.22795 | |
| 14 | | 0.000557 | | | 193 | | 0.146 | | | 0.143 | | -114.63312 | | | 60.7244 | |
| 15 | | 0.000535 | | | 144 | | 0.180 | | | 0.286 | | -116.4269 | | | 60.45934 | |
| 16 | | 0.000568 | | | 269 | | 0.204 | | | 0.306 | | -116.54705 | | | 60.01954 | |
| 17 | | 0.000581 | | | 424 | | 0.186 | | | 0.200 | | -114.77958 | | | 59.93499 | |
| 18 | | 0.000502 | | | 24 | | 0.184 | | | 0.167 | | -113.777 | | | 59.67629 | |
| 19 | | 0.000639 | | | 270 | | 0.223 | | | 0.067 | | -103.6514 | | | 54.24964 | |
| 20 | | 0.000494 | | | 66 | | 0.161 | | | 0.267 | | -112.67055 | | | 57.81732 | |
| 21 | | 0.000513 | | | 49 | | 0.155 | | | 0.000 | | -103.52096 | | | 55.60701 | |
| 22 | | 0.000659 | | | 218 | | 0.307 | | | 0.200 | | -102.03099 | | | 55.08039 | |
| 23 | | 0.000659 | | | 209 | | 0.197 | | | 0.111 | | -95.739344 | | | 53.07447 | |
| 24 | | 0.000627 | | | 39 | | 0.287 | | | 0.267 | | -95.478835 | | | 56.38909 | |
| 25 | | 0.000696 | | | 299 | | 0.293 | | | 0.095 | | -93.774004 | | | 56.25846 | |
| 26 | | 0.000625 | | | 76 | | 0.253 | | | 0.100 | | -97.297339 | | | 53.92717 | |
| 27 | | 0.000652 | | | 211 | | 0.299 | | | 0.238 | | -90.049586 | | | 56.73238 | |
| 28 | | 0.000484 | | | 30 | | 0.176 | | | 0.000 | | -96.339726 | | | 52.05528 | |
| 29 | | 0.000435 | | | 2 | | 0.166 | | | 0.333 | | -96.977681 | | | 52.60875 | |
| 30 | | 0.000595 | | | 114 | | 0.167 | | | 0.200 | | -95.841789 | | | 51.42894 | |
| 31 | | 0.000603 | | | 34 | | 0.200 | | | 0.214 | | -95.229662 | | | 50.77985 | |
| 32 | | 0.000397 | | | 0 | | 0.186 | | | 0.833 | | -126.69331 | | | 65.44044 | |
| 33 | | 0.000414 | | | 13 | | 0.175 | | | 0.500 | | -129.09502 | | | 66.62309 | |
| 34 | | 0.000443 | | | 50 | | 0.092 | | | 0.267 | | -128.48412 | | | 66.25545 | |
| 35 | | 0.000419 | | | 32 | | 0.211 | | | 0.667 | | -125.25634 | | | 65.05612 | |
| 36 | | 0.000678 | | | 226 | | 0.225 | | | 0.143 | | -105.70477 | | | 55.29422 | |
| 37 | | 0.00062 | | | 91 | | 0.241 | | | 0.200 | | -104.28357 | | | 54.99741 | |
| 38 | | 0.000503 | | | 141 | | 0.190 | | | 0.200 | | -122.56139 | | | 61.93277 | |
| 39 | | 0.000516 | | | 67 | | 0.194 | | | 0.200 | | -109.23527 | | | 56.36965 | |
| 40 | | 0.000583 | | | 262 | | 0.186 | | | 0.167 | | -109.94397 | | | 55.82088 | |
| 41 | | 0.000463 | | | 6 | | 0.146 | | | 0.300 | | -112.66548 | | | 55.71534 | |
| 42 | | 0.00053 | | | 37 | | 0.185 | | | 0.333 | | -111.23483 | | | 55.52859 | |
| 43 | | 0.000337 | | | 0 | | 0.061 | | | 0.095 | | -111.12941 | | | 56.61858 | |
| 44 | | 0.000613 | | | 51 | | 0.226 | | | 0.133 | | -93.188295 | | | 56.64026 | |
| 45 | | 0.000616 | | | 96 | | 0.190 | | | 0.067 | | -95.675878 | | | 55.74407 | |
| 46 | | 0.000529 | | | 2 | | 0.176 | | | 0.333 | | -92.367535 | | | 51.10916 | |
| 47 | | 0.000593 | | | 120 | | 0.199 | | | 0.278 | | -92.028 | | | 51.63134 | |
| 48 | | 0.000552 | | | 35 | | 0.153 | | | 0.214 | | -87.013668 | | | 48.64702 | |
| 49 | | 0.000604 | | | 49 | | 0.212 | | | 0.286 | | -83.895104 | | | 53.73871 | |
| 50 | | 0.000547 | | | 31 | | 0.221 | | | 0.300 | | -84.92319 | | | 52.68658 | |
| 51 | | 0.00043 | | | 11 | | 0.108 | | | 0.000 | | -83.636956 | | | 52.13778 | |
| 52 | | 0.000642 | | | 125 | | 0.212 | | | 0.273 | | -84.939411 | | | 51.28054 | |
| 53 | | 0.000666 | | | 163 | | 0.216 | | | 0.139 | | -86.510032 | | | 52.01633 | |
| 54 | | 0.000536 | | | 42 | | 0.148 | | | 0.286 | | -83.302452 | | | 50.79985 | |
| 55 | | 0.000468 | | | 8 | | 0.159 | | | 0.333 | | -81.894188 | | | 50.10322 | |
| 56 | | 0.000547 | | | 16 | | 0.153 | | | 0.333 | | -82.404066 | | | 51.8241 | |
| 57 | | 0.000654 | | | 184 | | 0.213 | | | 0.256 | | -90.106001 | | | 52.78016 | |
| 58 | | 0.000628 | | | 79 | | 0.189 | | | 0.200 | | -87.16348 | | | 53.26777 | |
| 59 | | 0.000536 | | | 74 | | 0.155 | | | 0.267 | | -92.400949 | | | 52.92082 | |
| 60 | | 0.000507 | | | 25 | | 0.149 | | | 0.300 | | -84.185374 | | | 51.05756 | |
| 61 | | 0.000611 | | | 68 | | 0.195 | | | 0.250 | | -89.378808 | | | 54.07371 | |
| 62 | | 0.000646 | | | 237 | | 0.157 | | | 0.214 | | -80.037376 | | | 49.96553 | |
| 63 | | 0.000646 | | | 238 | | 0.189 | | | 0.071 | | -89.380661 | | | 55.55084 | |
| 64 | | 0.000647 | | | 58 | | 0.233 | | | 0.179 | | -90.065591 | | | 50.31271 | |
| 65 | | 0.000613 | | | 36 | | 0.247 | | | 0.429 | | -94.131722 | | | 51.91145 | |
| 66 | | 0.000664 | | | 258 | | 0.229 | | | 0.267 | | -91.328846 | | | 51.15917 | |
| 67 | | 0.000611 | | | 213 | | 0.182 | | | 0.200 | | -86.944026 | | | 50.15652 | |
| 68 | | 0.000675 | | | 264 | | 0.217 | | | 0.200 | | -87.907256 | | | 50.87773 | |
| 69 | | 0.00063 | | | 75 | | 0.215 | | | 0.133 | | -90.233415 | | | 51.49843 | |
| 70 | | 0.000506 | | | 63 | | 0.176 | | | 0.300 | | -110.42884 | | | 55.07268 | |
| 71 | | 0.000429 | | | 9 | | 0.145 | | | 0.500 | | -113.9908 | | | 56.23017 | |
| 72 | | 0.000529 | | | 134 | | 0.186 | | | 0.250 | | -112.79645 | | | 56.74931 | |
| 73 | | 0.000525 | | | 135 | | 0.172 | | | 0.286 | | -114.49502 | | | 57.48293 | |
| 74 | | 0.000594 | | | 34 | | 0.262 | | | 0.333 | | -102.85791 | | | 54.51913 | |
| 75 | | 0.000466 | | | 136 | | 0.185 | | | 0.400 | | -124.00692 | | | 65.53797 | |
| 76 | | 0.000633 | | | 268 | | 0.160 | | | 0.100 | | -103.7306 | | | 57.39444 | |
| 77 | | 0.000674 | | | 285 | | 0.147 | | | 0.121 | | -103.77208 | | | 56.48986 | |
| 78 | | 0.00062 | | | 216 | | 0.213 | | | 0.333 | | -107.47442 | | | 57.29227 | |
| 79 | | 0.000628 | | | 145 | | 0.177 | | | 0.333 | | -105.16368 | | | 57.40266 | |
| 80 | | 0.000657 | | | 256 | | 0.200 | | | 0.357 | | -106.19356 | | | 56.53577 | |
| 81 | | 0.000625 | | | 264 | | 0.266 | | | 0.167 | | -107.50183 | | | 55.35125 | |
| 82 | | 0.00058 | | | 213 | | 0.223 | | | 0.333 | | -108.23413 | | | 55.60412 | |
| 83 | | 0.000435 | | | 34 | | 0.144 | | | 0.167 | | -116.79455 | | | 61.06033 | |
| 84 | | 0.000578 | | | 412 | | 0.186 | | | 0.273 | | -117.96365 | | | 61.07467 | |
| 85 | | 0.000534 | | | 97 | | 0.158 | | | 0.267 | | -119.12258 | | | 59.94259 | |
| 86 | | 0.000511 | | | 25 | | 0.183 | | | 0.476 | | -118.62766 | | | 60.67314 | |
| 87 | | 0.000476 | | | 9 | | 0.177 | | | 0.200 | | -117.96732 | | | 59.58456 | |
| 88 | | 0.00053 | | | 206 | | 0.142 | | | 0.133 | | -120.3421 | | | 61.41613 | |
| 89 | | 0.000536 | | | 90 | | 0.154 | | | 0.467 | | -122.26318 | | | 60.25524 | |
| 90 | | 0.000404 | | | 9 | | 0.092 | | | 0.167 | | -116.56213 | | | 62.24475 | |
| 91 | | 0.000376 | | | 4 | | 0.136 | | | 0.000 | | -116.39623 | | | 61.70466 | |
| 92 | | 0.0005 | | | 73 | | 0.122 | | | 0.133 | | -115.24343 | | | 56.58098 | |
| 93 | | 0.000564 | | | 130 | | 0.173 | | | 0.200 | | -109.50644 | | | 57.8307 | |
| 94 | | 0.000416 | | | 33 | | 0.132 | | | 0.333 | | -114.51654 | | | 55.29706 | |
| 95 | | 0.000436 | | | 23 | | 0.150 | | | 0.400 | | -114.94073 | | | 55.84953 | |
| 96 | | 0.000502 | | | 42 | | 0.148 | | | 0.200 | | -110.55336 | | | 57.22509 | |
| 97 | | 0.000504 | | | 24 | | 0.141 | | | 0.133 | | -110.20652 | | | 58.54929 | |
| 98 | | 0.000532 | | | 62 | | 0.196 | | | 0.200 | | -110.64909 | | | 57.89768 | |
| 99 | | 0.000574 | | | 21 | | 0.204 | | | 0.400 | | -108.31324 | | | 56.835 | |
| 100 | | 0.00055 | | | 60 | | 0.122 | | | 0.200 | | -105.43252 | | | 56.20399 | |
| 101 | | 0.000613 | | | 156 | | 0.231 | | | 0.600 | | -107.37825 | | | 56.64872 | |
| 102 | | 0.000608 | | | 90 | | 0.182 | | | 0.143 | | -95.483058 | | | 56.13588 | |
| 103 | | 0.000509 | | | 5 | | 0.140 | | | 0.400 | | -85.75638 | | | 47.70972 | |
| Table S2. Summary statistics calculated for nodes after sample size was adjusted to n=15 for all nodes as per methods described in main manuscript. Sample size (N), number of alleles (Na), number of effective alleles (Ne), Shannon’s Information Index (I), observed heterozygosity (Ho), expected heterozygosity (He), unbiased expected heterozygosity (uHe), and fixation index (F). Summary statistics were calculated using GenAlEx 6.5. | | | | | | | | | | | | | | | | |
| Node | **N** | | **Na** | **Ne** | | **I** | | **Ho** | **He** | | **uHe** | | **F** | **Latitude** | | **Longitude** |
| 1 | 15 | | 5.00 | 2.97 | | 1.18 | | 0.58 | 0.59 | | 0.61 | | 0.03 | 52.95832 | | -98.9544 |
| 2 | 15 | | 5.56 | 3.26 | | 1.33 | | 0.74 | 0.65 | | 0.68 | | -0.13 | 53.58931 | | -101.237 |
| 3 | 15 | | 6.22 | 3.58 | | 1.46 | | 0.72 | 0.69 | | 0.72 | | -0.04 | 54.91077 | | -101.408 |
| 4 | 15 | | 7.00 | 4.13 | | 1.58 | | 0.72 | 0.72 | | 0.75 | | -0.01 | 54.17651 | | -99.5473 |
| 5 | 15 | | 6.89 | 3.35 | | 1.49 | | 0.66 | 0.70 | | 0.72 | | 0.04 | 54.57689 | | -105.362 |
| 6 | 15 | | 6.67 | 3.97 | | 1.53 | | 0.68 | 0.71 | | 0.74 | | 0.03 | 54.75783 | | -105.84 |
| 7 | 15 | | 6.22 | 3.17 | | 1.33 | | 0.59 | 0.62 | | 0.64 | | 0.06 | 53.96669 | | -105.814 |
| 8 | 15 | | 5.56 | 3.31 | | 1.35 | | 0.72 | 0.67 | | 0.70 | | -0.04 | 54.45689 | | -106.617 |
| 9 | 15 | | 6.11 | 3.79 | | 1.47 | | 0.76 | 0.70 | | 0.73 | | -0.06 | 54.35277 | | -107.198 |
| 10 | 15 | | 6.67 | 3.93 | | 1.49 | | 0.71 | 0.70 | | 0.72 | | 0.00 | 54.52597 | | -100.174 |
| 11 | 15 | | 3.67 | 2.50 | | 1.03 | | 0.62 | 0.58 | | 0.60 | | -0.07 | 52.1309 | | -98.2955 |
| 12 | 15 | | 6.00 | 3.76 | | 1.43 | | 0.70 | 0.69 | | 0.71 | | -0.03 | 55.2376 | | -100.321 |
| 13 | 15 | | 6.33 | 3.72 | | 1.48 | | 0.69 | 0.71 | | 0.74 | | 0.05 | 55.05928 | | -98.7064 |
| 14 | 15 | | 6.56 | 4.18 | | 1.58 | | 0.74 | 0.74 | | 0.77 | | 0.00 | 60.86263 | | -114.778 |
| 15 | 15 | | 7.33 | 4.03 | | 1.60 | | 0.76 | 0.74 | | 0.76 | | -0.04 | 60.55704 | | -116.366 |
| 16 | 15 | | 7.22 | 4.63 | | 1.67 | | 0.73 | 0.76 | | 0.79 | | 0.03 | 60.19195 | | -116.841 |
| 17 | 15 | | 7.00 | 4.45 | | 1.63 | | 0.75 | 0.75 | | 0.78 | | 0.00 | 59.92033 | | -114.831 |
| 18 | 15 | | 7.00 | 4.16 | | 1.61 | | 0.76 | 0.74 | | 0.77 | | -0.03 | 59.77583 | | -113.785 |
| 19 | 15 | | 6.22 | 4.06 | | 1.52 | | 0.70 | 0.73 | | 0.76 | | 0.04 | 54.31684 | | -103.107 |
| 20 | 15 | | 6.67 | 4.39 | | 1.60 | | 0.78 | 0.75 | | 0.77 | | -0.04 | 58.01268 | | -112.989 |
| 21 | 15 | | 5.11 | 3.21 | | 1.28 | | 0.62 | 0.64 | | 0.67 | | 0.06 | 55.49964 | | -103.691 |
| 22 | 15 | | 6.67 | 3.80 | | 1.49 | | 0.62 | 0.68 | | 0.71 | | 0.11 | 54.8406 | | -102.273 |
| 23 | 15 | | 6.56 | 3.25 | | 1.41 | | 0.65 | 0.67 | | 0.69 | | 0.02 | 53.0606 | | -95.7486 |
| 24 | 15 | | 5.78 | 3.38 | | 1.37 | | 0.70 | 0.68 | | 0.71 | | -0.02 | 56.3847 | | -95.4965 |
| 25 | 15 | | 6.11 | 3.49 | | 1.41 | | 0.66 | 0.68 | | 0.70 | | 0.02 | 56.36068 | | -93.7819 |
| 26 | 15 | | 6.00 | 3.59 | | 1.43 | | 0.68 | 0.70 | | 0.72 | | 0.02 | 53.95209 | | -97.0653 |
| 27 | 15 | | 6.44 | 3.47 | | 1.43 | | 0.67 | 0.68 | | 0.70 | | 0.03 | 56.736 | | -90.0509 |
| 28 | 15 | | 5.89 | 3.24 | | 1.35 | | 0.72 | 0.66 | | 0.69 | | -0.10 | 51.99492 | | -96.2229 |
| 29 | 15 | | 5.89 | 3.41 | | 1.38 | | 0.61 | 0.67 | | 0.69 | | 0.09 | 51.32417 | | -95.875 |
| 30 | 15 | | 5.44 | 2.99 | | 1.28 | | 0.67 | 0.65 | | 0.67 | | -0.04 | 50.822 | | -94.4892 |
| 31 | 15 | | 7.89 | 5.01 | | 1.77 | | 0.84 | 0.78 | | 0.81 | | -0.07 | 65.26 | | -126.53 |
| 32 | 15 | | 7.11 | 4.60 | | 1.67 | | 0.77 | 0.76 | | 0.79 | | -0.01 | 66.85305 | | -128.831 |
| 33 | 15 | | 6.89 | 4.33 | | 1.63 | | 0.76 | 0.75 | | 0.78 | | -0.02 | 65.03472 | | -125.462 |
| 34 | 15 | | 6.44 | 3.67 | | 1.48 | | 0.67 | 0.71 | | 0.73 | | 0.06 | 55.21178 | | -105.431 |
| 35 | 15 | | 6.11 | 3.77 | | 1.46 | | 0.72 | 0.70 | | 0.73 | | -0.02 | 55.06511 | | -104.633 |
| 36 | 15 | | 7.44 | 5.01 | | 1.75 | | 0.80 | 0.79 | | 0.82 | | -0.02 | 62.42071 | | -121.88 |
| 37 | 15 | | 6.22 | 3.56 | | 1.45 | | 0.72 | 0.70 | | 0.73 | | -0.02 | 56.29732 | | -109.714 |
| 38 | 15 | | 6.22 | 3.53 | | 1.42 | | 0.66 | 0.68 | | 0.70 | | 0.07 | 56.02578 | | -110.461 |
| 39 | 15 | | 5.56 | 3.48 | | 1.39 | | 0.58 | 0.69 | | 0.72 | | 0.16 | 55.9256 | | -112.87 |
| 40 | 15 | | 7.11 | 4.15 | | 1.62 | | 0.79 | 0.75 | | 0.78 | | -0.06 | 55.40771 | | -111.186 |
| 41 | 15 | | 6.11 | 3.47 | | 1.41 | | 0.69 | 0.68 | | 0.71 | | -0.02 | 56.46687 | | -93.415 |
| 42 | 15 | | 6.44 | 3.56 | | 1.48 | | 0.64 | 0.70 | | 0.73 | | 0.08 | 56.21349 | | -95.5479 |
| 43 | 15 | | 5.56 | 3.26 | | 1.26 | | 0.68 | 0.61 | | 0.64 | | -0.11 | 51.20188 | | -92.3352 |
| 44 | 15 | | 6.11 | 3.23 | | 1.36 | | 0.61 | 0.65 | | 0.68 | | 0.08 | 51.25733 | | -92.5957 |
| 45 | 15 | | 3.56 | 2.64 | | 1.04 | | 0.62 | 0.58 | | 0.61 | | -0.06 | 48.64823 | | -86.9631 |
| 46 | 15 | | 6.56 | 3.72 | | 1.49 | | 0.69 | 0.70 | | 0.73 | | 0.02 | 52.71339 | | -85.0705 |
| 47 | 15 | | 6.33 | 3.38 | | 1.43 | | 0.66 | 0.68 | | 0.71 | | 0.02 | 52.15119 | | -83.0795 |
| 48 | 15 | | 6.11 | 3.03 | | 1.36 | | 0.62 | 0.66 | | 0.68 | | 0.06 | 51.14038 | | -84.1996 |
| 49 | 15 | | 6.00 | 3.03 | | 1.33 | | 0.58 | 0.64 | | 0.66 | | 0.09 | 51.71963 | | -86.0471 |
| 50 | 15 | | 6.44 | 3.67 | | 1.49 | | 0.78 | 0.72 | | 0.75 | | -0.07 | 50.93843 | | -83.4053 |
| 51 | 15 | | 6.00 | 3.59 | | 1.44 | | 0.69 | 0.69 | | 0.72 | | 0.01 | 50.54864 | | -81.9165 |
| 52 | 15 | | 6.11 | 3.32 | | 1.40 | | 0.63 | 0.68 | | 0.71 | | 0.08 | 52.46398 | | -90.6205 |
| 53 | 15 | | 6.33 | 3.24 | | 1.40 | | 0.70 | 0.67 | | 0.70 | | -0.04 | 53.85379 | | -89.0416 |
| 54 | 15 | | 5.56 | 3.47 | | 1.41 | | 0.73 | 0.70 | | 0.72 | | -0.05 | 50.0143 | | -79.9969 |
| 55 | 15 | | 5.78 | 3.32 | | 1.36 | | 0.63 | 0.67 | | 0.70 | | 0.07 | 55.76913 | | -89.1141 |
| 56 | 15 | | 5.89 | 3.06 | | 1.31 | | 0.65 | 0.64 | | 0.66 | | -0.02 | 50.59366 | | -90.514 |
| 57 | 15 | | 6.44 | 3.56 | | 1.45 | | 0.62 | 0.70 | | 0.73 | | 0.13 | 51.90233 | | -94.2725 |
| 58 | 15 | | 6.33 | 3.10 | | 1.34 | | 0.63 | 0.65 | | 0.67 | | 0.04 | 51.22389 | | -91.1527 |
| 59 | 15 | | 6.11 | 3.41 | | 1.42 | | 0.66 | 0.68 | | 0.70 | | 0.04 | 50.25328 | | -85.4437 |
| 60 | 15 | | 5.89 | 2.98 | | 1.33 | | 0.68 | 0.65 | | 0.67 | | -0.05 | 50.70833 | | -87.6833 |
| 61 | 15 | | 5.56 | 2.96 | | 1.30 | | 0.63 | 0.65 | | 0.67 | | 0.00 | 51.56593 | | -90.5069 |
| 62 | 15 | | 7.00 | 4.19 | | 1.62 | | 0.81 | 0.75 | | 0.77 | | -0.08 | 55.14513 | | -110.136 |
| 63 | 15 | | 7.00 | 4.26 | | 1.61 | | 0.77 | 0.75 | | 0.77 | | -0.03 | 56.2276 | | -114.3 |
| 64 | 15 | | 6.33 | 3.87 | | 1.48 | | 0.69 | 0.70 | | 0.72 | | 0.00 | 56.71633 | | -111.999 |
| 65 | 15 | | 6.78 | 4.38 | | 1.63 | | 0.70 | 0.76 | | 0.79 | | 0.08 | 57.35268 | | -114.722 |
| 66 | 15 | | 6.67 | 3.91 | | 1.55 | | 0.70 | 0.73 | | 0.76 | | 0.05 | 54.47193 | | -103.086 |
| 67 | 15 | | 7.56 | 4.81 | | 1.71 | | 0.83 | 0.77 | | 0.80 | | -0.08 | 65.53183 | | -123.664 |
| 68 | 15 | | 6.56 | 3.91 | | 1.49 | | 0.69 | 0.70 | | 0.72 | | 0.01 | 56.83132 | | -103.433 |
| 69 | 15 | | 7.67 | 4.67 | | 1.70 | | 0.81 | 0.77 | | 0.80 | | -0.06 | 57.16457 | | -107.246 |
| 70 | 15 | | 7.22 | 4.33 | | 1.63 | | 0.75 | 0.75 | | 0.78 | | 0.00 | 57.23338 | | -105.524 |
| 71 | 15 | | 7.22 | 4.61 | | 1.61 | | 0.78 | 0.73 | | 0.75 | | -0.07 | 56.54674 | | -105.771 |
| 72 | 15 | | 6.89 | 4.03 | | 1.54 | | 0.74 | 0.72 | | 0.74 | | -0.04 | 55.3855 | | -107.336 |
| 73 | 15 | | 6.33 | 3.91 | | 1.53 | | 0.69 | 0.74 | | 0.76 | | 0.07 | 55.35349 | | -108.406 |
| 74 | 15 | | 7.33 | 4.39 | | 1.66 | | 0.73 | 0.76 | | 0.79 | | 0.04 | 60.9465 | | -116.55 |
| 75 | 15 | | 7.56 | 4.76 | | 1.70 | | 0.74 | 0.76 | | 0.79 | | 0.01 | 61.09321 | | -118.073 |
| 76 | 15 | | 8.00 | 4.76 | | 1.75 | | 0.77 | 0.77 | | 0.80 | | 0.01 | 60.03347 | | -118.826 |
| 77 | 15 | | 7.56 | 4.57 | | 1.68 | | 0.79 | 0.75 | | 0.78 | | -0.04 | 60.6535 | | -118.518 |
| 78 | 15 | | 7.56 | 4.43 | | 1.68 | | 0.77 | 0.76 | | 0.79 | | -0.01 | 60.0218 | | -117.614 |
| 79 | 15 | | 7.33 | 4.46 | | 1.66 | | 0.81 | 0.76 | | 0.78 | | -0.08 | 61.40061 | | -120.951 |
| 80 | 15 | | 7.78 | 4.60 | | 1.73 | | 0.76 | 0.78 | | 0.81 | | 0.03 | 62.27182 | | -116.316 |
| 81 | 15 | | 7.00 | 4.41 | | 1.62 | | 0.76 | 0.75 | | 0.78 | | -0.02 | 61.86537 | | -116.337 |
| 82 | 15 | | 5.89 | 3.76 | | 1.47 | | 0.69 | 0.71 | | 0.74 | | 0.04 | 56.37232 | | -115.174 |
| 83 | 15 | | 7.78 | 4.29 | | 1.67 | | 0.80 | 0.75 | | 0.78 | | -0.05 | 57.58481 | | -109.734 |
| 84 | 15 | | 4.33 | 2.93 | | 1.17 | | 0.66 | 0.63 | | 0.65 | | -0.06 | 55.30024 | | -114.678 |
| 85 | 15 | | 4.67 | 3.10 | | 1.24 | | 0.74 | 0.66 | | 0.68 | | -0.12 | 55.7974 | | -114.842 |
| 86 | 15 | | 5.89 | 3.61 | | 1.42 | | 0.68 | 0.69 | | 0.72 | | 0.03 | 57.21673 | | -110.647 |
| 87 | 15 | | 7.33 | 4.86 | | 1.72 | | 0.89 | 0.78 | | 0.81 | | -0.14 | 57.80435 | | -110.718 |
| 88 | 15 | | 7.67 | 5.08 | | 1.75 | | 0.83 | 0.78 | | 0.81 | | -0.06 | 56.77202 | | -108.059 |
| 89 | 15 | | 5.56 | 3.79 | | 1.45 | | 0.76 | 0.72 | | 0.74 | | -0.06 | 56.09928 | | -105.611 |
| 90 | 15 | | 7.11 | 3.77 | | 1.51 | | 0.68 | 0.69 | | 0.71 | | 0.02 | 56.49043 | | -107.568 |
